# 3D Printed X-ray Compatible Microfluidics for Online Characterization of Hexosomes: A Synchrotron SAXS-on-Chip Study with Molecular Dynamics Insights

**DOI:** 10.64898/2026.08.31.748233

**Authors:** Zahra Babaie, Mariana Valério, Fabian Schuhmann, Maria Dimaki, Babak Rezaei, Weria Pezeshkian, Stephan S. Keller, Winnie E. Svendsen, Paulo C. T. Souza, Anan Yaghmur

**Author notes:** Corresponding Author Anan Yaghmur —. If an author’s address is different than the one given in the affiliation line, this information may be included here (Anan Yaghmur).

## Abstract

Online structural characterization during microfluidic lipid self-assembly is important for understanding and controlling the formation of nonlamellar liquid crystalline nanodispersions. Here, we report a 3D-printed, X-ray-compatible hydrodynamic flow-focusing microfluidic chip with variable channel dimensions, integrated with synchrotron small-angle X-ray scattering (SAXS), for position-resolved SAXS-on-chip monitoring of Ca^2+^-triggered hexosome formation. Hexosomes were produced under continuous flow by mixing ethanolic solutions of docosahexaenoic acid monoglyceride (MAG-DHA), the negatively charged phosphatidylglycerol DOPG, and α-tocopherol with Ca^2+^-containing PIPES buffer. Online SAXS-on-chip measurements detected three Bragg reflections characteristic of the internal inverse hexagonal (H_2_) phase on a tens-of-milliseconds residence-time scale, revealing rapid structural evolution during microfluidic mixing. Complementary *ex situ* SAXS identified the DOPG/Ca^2+^ molar ratio as a key parameter modulating the direct vesicle-to-hexosome transformation and the compactness of the internal H_2_ nanostructures. Dynamic light scattering showed that the flow-rate ratio modulated nanoparticle size, yielding hexosomes with mean hydrodynamic diameters in the range of approximately 120-175 nm and polydispersity index values down to 0.14 at a total flow rate of 200 µL min^−1^. Cryo-TEM revealed coexistence of hexosomes and vesicular nanostructures, highlighting morphological heterogeneity, while Coarse-Grained Molecular Dynamics simulations supported a central role of Ca^2+^-DOPG association in promoting a direct lamellar-H_2_ phase transition. Overall, this work shows that 3D-printed SAXS-compatible microfluidics can integrate continuous production with online structural characterization, providing a basis for future formulation and process optimization of drug-loaded cubosomes, hexosomes, and related nonlamellar liquid crystalline nanodispersions.

## 1. INTRODUCTION

Microfluidics offers unprecedented opportunities to advance research in drug delivery, diagnostics, and biomedicine^1,2^. Through precise control of the flow dynamics, small volumes, and mixing conditions, microfluidics has also revolutionized the synthesis of nanomaterials with controllable sizes and shapes and potential production scalability^3,4^. In recent years, there has been rapid growth in the fabrication of two-dimensional (2D) and three-dimensional (3D) microfluidic platforms for the synthesis of soft and inorganic nanomaterials for various applications^5,6^. Further, they have a great potential to advance research on nanoparticle production kinetics and structural dynamics under confined geometry conditions through coupling specially designed X-ray (or neutron) compatible microfluidic platforms with synchrotron small-angle X-ray (SAXS) or neutron (SANS) scattering techniques^5,7–9^.

Designing X-ray compatible microfluidic chips is particularly demanding due to a range of interconnected challenges. Achieving high X-ray transmission, for instance, requires thin materials composed of low atomic number elements (Z ≤ 14, where Z is the atomic number)^10^. At the same time, resistance to prolonged X-ray exposure necessitates the use of specialized materials such as Kapton foils^11–15^, cyclic olefin copolymers^16–19^, or alternatively the incorporation of dedicated X-ray transparent windows into microfluidic devices^20,21^. Minimizing X-ray attenuation and background scattering is another critical consideration, as it directly affects data reliability by maintaining a high signal-to-noise ratio^22^. Furthermore, reproducibility across microfluidic platforms depends on fabrication precision, particularly the production of consistent channel dimensions that remain stable throughout the experiment^5,13^. Optical transparency is an essential requirement, enabling real-time optical monitoring of fluid flow within the microchannels and facilitating accurate alignment of the device with the incident X-rays^10^. Notably, similar design constraints, including beam attenuation minimization and background scattering control, are encountered in designing microfluidics for SANS investigations^23,24^.

From an experimental robustness standpoint, practical considerations such as the chemical compatibility of X-ray compatible microfluidic chips with organic solvents (typically ethanol), ease of cleaning, and durability for repeated experiments are critical for microfluidic chip design^5,12,25^. Preventing adsorption of lipids and biomolecules on microchannel walls is also a critical consideration, as such surface fouling can lead to clogging and compromise X-ray measurement quality. Surface fouling can be avoided through microfluidic surface modification strategies, including PEGylation, oxygen plasma treatment, and the application of hydrophilic polymer coatings^26–28^.

Despite the attractiveness and high potential of microfluidic devices in generating lamellar (vesicles)^29^ and non-lamellar nano-self-assemblies (such as cubosomes and hexosomes) ^30–35^, only a few studies reported on online SAXS-on-chip investigations for real-time monitoring of the nanoparticle formation kinetics and involved self-assembly dynamics during their microfluidic synthesis^5,15,30^. In these few studies, the SAXS-on-chip experiments were conducted using 2D hydrodynamic flow-focusing (HFF) microfluidic chips with relatively wide channels (∼300 µm) to avoid parasitic scattering from the channel walls and guarantee a successful SAXS analysis through achieving a good signal-to-noise ratio. However, these relatively wider channels compromise the microfluidic efficacy in controlling the NP size characteristics^5,15^. To overcome these limitations, we introduce, for the first time, 3D-printed X-ray compatible microfluidic chip, featuring a serpentine channel with alternating widths of 100 µm and 270 µm. The narrower segments (100 µm) are designed to enhance mixing kinetics and size control by reducing diffusion distances, while the wider zones (270 µm) serve as observation windows, optimized to increase the X-ray path length, thereby improving the signal-to-noise ratio and enabling successful SAXS analysis.

Several, 3D HFF microfluidic designs have been developed to position the focused stream at the center of the 3D parabolic flow profile, where velocity gradients are minimized, resulting in more uniform mixing compared to planar geometries^13,31^. However, the fabrication of 3D focusing geometries is complex, as conventional approaches such as multilayer soft lithography ^32,33^ and Kapton-based microfabrication^13^ require multiple alignments and bonding steps, which limit device architecture and flow throughput. In this context, 3D printing has emerged in recent years as a versatile strategy that minimizes fabrication time and cost for the rapid prototyping of monolithic microfluidic devices. It enables the integration of complex microfluidic features within compact architectures^34,35^.

In this study, we report on 3D-printing of X-ray compatible microfluidic chips with variable dimensions and its combination with synchrotron SAXS for online structural analysis of hexosomes during their continuous production. The nanodispersions are continuously produced through spontaneous self-assembly of a ternary lipid mixture, consisting of docosahexaenoic acid monoacylglycerol (MAG-DHA), phosphatidylglycerol (1,2-dioleoyl-sn-glycero-3-phosphoglycerol, DOPG), and α-tocopherol, in buffer environment (PIPES buffer, 10 mM, pH 7.0), containing Ca^2+^ ions and the polymeric stabilizer Pluronic F127. It is worth mentioning that MAG-DHA is an amphiphilic bioactive lipid synthesized through sn-1 esterification of glycerol with ethyl docosahexaenoate, and attractive as a metabolizable precursor to docosahexaenoic acid (DHA; 22:6, n-3), a biologically important omega-3 fatty acid known for its neuroprotective and anti-inflammatory properties^36,37^. Our SAXS experimental findings revealed rapid hexosome formation (≥14 ms) upon exposure of the ethanolic solution of the ternary lipid system to a buffer containing Ca^2+^ ions within a microfluidic channel. In addition to synchrotron SAXS, the produced nanodispersions were characterized using cryogenic transmission electron microscopy (cryo-TEM) and dynamic light scattering (DLS). This study demonstrates that the observed Ca^2+^-induced direct colloidal vesicle-hexosome transformation is associated with significant changes in the morphological features and size characteristics of the nanodispersions. The proposed mechanism underlying this transformation is further supported by molecular dynamics (MD) simulations.

## 2. EXPERIMENTAL SECTION

### 2.1 Materials

Docosahexaenoic acid monoacylglycerol (MAG-DHA) with a purity of >99% was purchased from Larodan (Solna, Sweden). 1,2-dioleoyl-sn-glycero-3-phospho-rac-glycerol, sodium salt (DOPG) was obtained from Lipoid GmbH (Ludwigshafen, Germany). D-α-tocopherol (α-vitamin E, α-Vit) with a purity of 98.28% was purchased from MedChemExpress (Monmouth Junction, NJ, USA). PIPES (1,4-piperazinediethanesulfonic acid, with a purity of 99%) and Pluronic^®^ F-127 were purchased from Sigma-Aldrich (St. Louis, MO, USA). The former was used to prepare 10 mM PIPES buffer at pH 7.0. Calcium chloride dihydrate (CaCl_2_·2H_2_O) and ethanol (EtOH) with a purity of 99% were purchased from VWR Chemicals (Søborg, Denmark). Ultrapure water was obtained from a Milli-Q system (Millipore Direct-Q3 ultraviolet system, MA, USA). All materials were used without further purification.

### 2.2. Microfluidic Chip Design and Fabrication

Recent advances in 3D printing have enabled rapid and precise fabrication of complex microfluidic devices^38^. However, digital light processing (DLP) desktop printers often suffer from unintended ultraviolet (UV) light exposure of trapped resin after the microchannel ceiling is printed, which can lead to partial polymerization and hinder subsequent resin removal^10^. To address this limitation and minimize exposure of trapped resin to stray light, high resolution projection micro-stereolithography (PµSL) technology combined with commercial resins designed for layer thicknesses < 20 µm can be employed. This enables the fabrication of monolithic microfluidic structures with thin suspended layers closing the microchannels eliminating the need for assembly and reducing the risk of leakage^39^. In addition, the high resolution of the PµSL printers allows the production of intricate microstructures with excellent dimensional accuracy and reproducibility^40^. Accordingly, PµSL technology was employed in this study to fabricate a 3D microfluidic chip, featuring a 40 mm-long serpentine channel with two alternating channel widths of 100 and 270 µm. A schematic illustration of the PµSL-based microfluidic chip fabrication process is shown in **Figure 1**. The wider channel sections (270 µm) were incorporated to facilitate *in situ* small-angle X-ray scattering (SAXS) measurements. This is important because reliable SAXS data analysis requires confining the X-ray beam between the microfluidic channel walls to avoid parasitic scattering from the channel surfaces^5,15^. The total resin thickness along the X-ray beam path was maintained at 400 µm, consisting of 200 µm layer of material both above and below the channel, to reduce X-ray attenuation and improve the signal-to-noise ratio during SAXS measurements.

**Figure 1.**
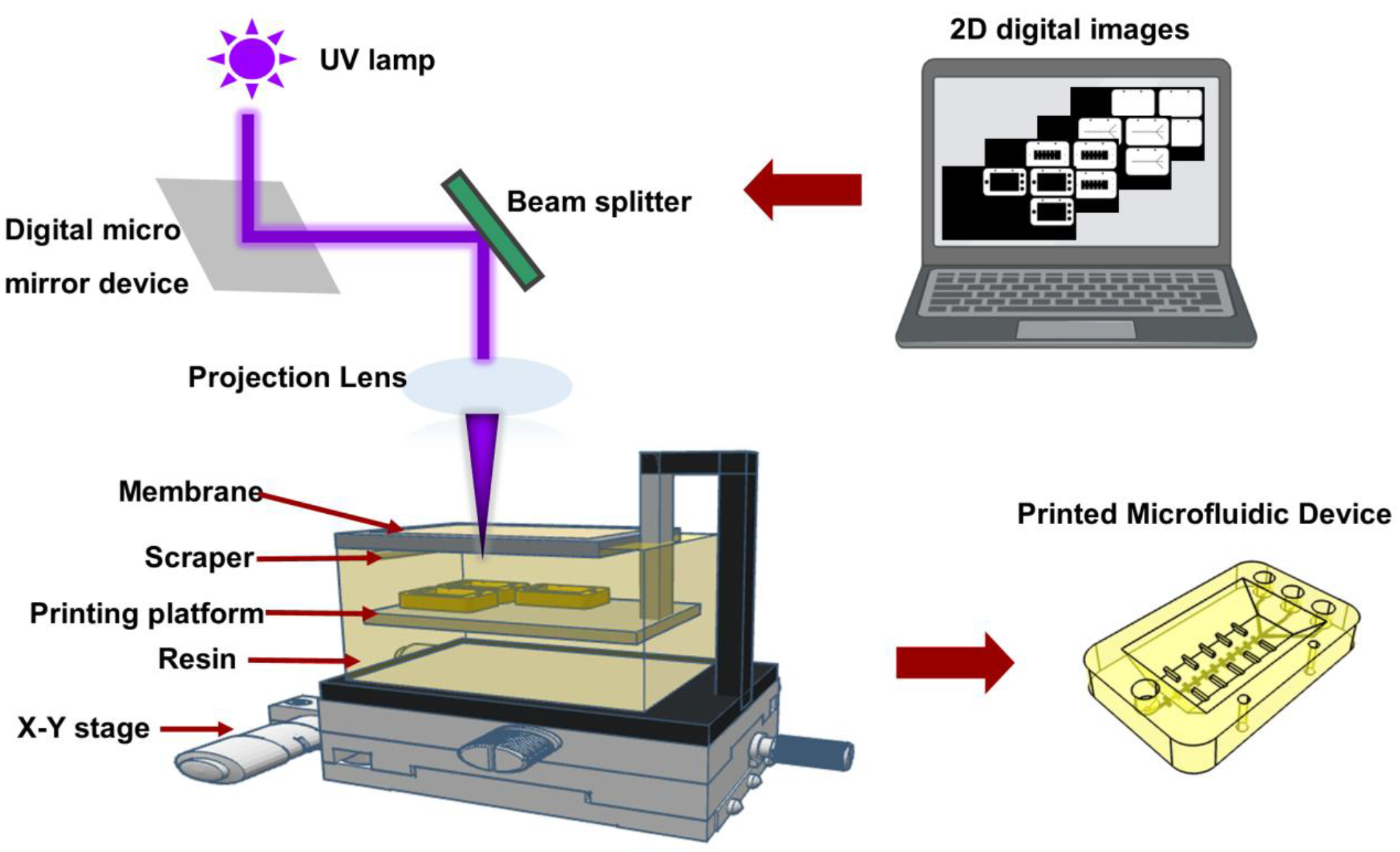
Schematic process flow for additive manufacturing of 3D microfluidic device.

Prior to fabrication, a 3D model of the microfluidic chip was designed using 3DEXPERIENCE CATIA (Dassault Systèmes, Vélizy-Villacoublay, France) and exported as standard tessellation language (STL) files. Detailed design parameters are presented in **Figure S1.** A hybrid layer-thickness strategy was applied during slicing by using Chitubox software (Chitubox Inc., Shenzhen, China). Specifically, the microchannel regions were sliced with a layer thickness of 5 µm to minimize wall roughness, while the remaining microstructural regions were sliced at 20 µm to reduce fabrication time without compromising device performance. Chip fabrication was carried out using a PµSL 3D printer (microArch® S140, BMF, MA, USA) and a commercially available high-temperature resin (HTL Yellow, 20 µm resolution; BMF, USA). Following printing, uncured resin was removed by rinsing the chip with isopropyl alcohol, followed by flushing the microchannels with pressurized nitrogen.

### 2.3. Continuous Production of Hexosomes

The continuous production of Pluronic F127-stabilized hexosomes was achieved through injection of PIPES buffer (10 mM, pH 7.0) containing 1.0 wt% Pluronic F127 and 5 mM Ca^2+^ into the two side channels (side inlets) of the chip, as illustrated in **Figure 2A**, using two 5 mL syringes (B. Braun, Melsungen, Germany). The two aqueous sheath streams hydrodynamically focused the central ethanol stream of ternary lipid mixtures introduced through the central microchannel (center inlet) by using a 1 mL Luer-lock syringe (Chirana, Stará Turá, Slovakia) both laterally and vertically and were subsequently mixed with the ethanol solution of lipids. In this study, two ethanol solutions of the ternary MAG-DHA/DOPG/α-tocopherol mixtures were prepared at a constant ethanol-to-lipid weight ratio of 60:40, with the following two lipid compositions: 48:40:12 and 56:30:14 (weight ratios), respectively. To enhance the limited solubility of DOPG in ethanol, an additional amount of water was added to the two ethanol solutions at final concentrations of 13.8 and 10.7 wt%, respectively, and the resulting solutions were placed in an incubator at 37 °C for 10 min before use to obtain clear stock solutions.

**Figure 2.**
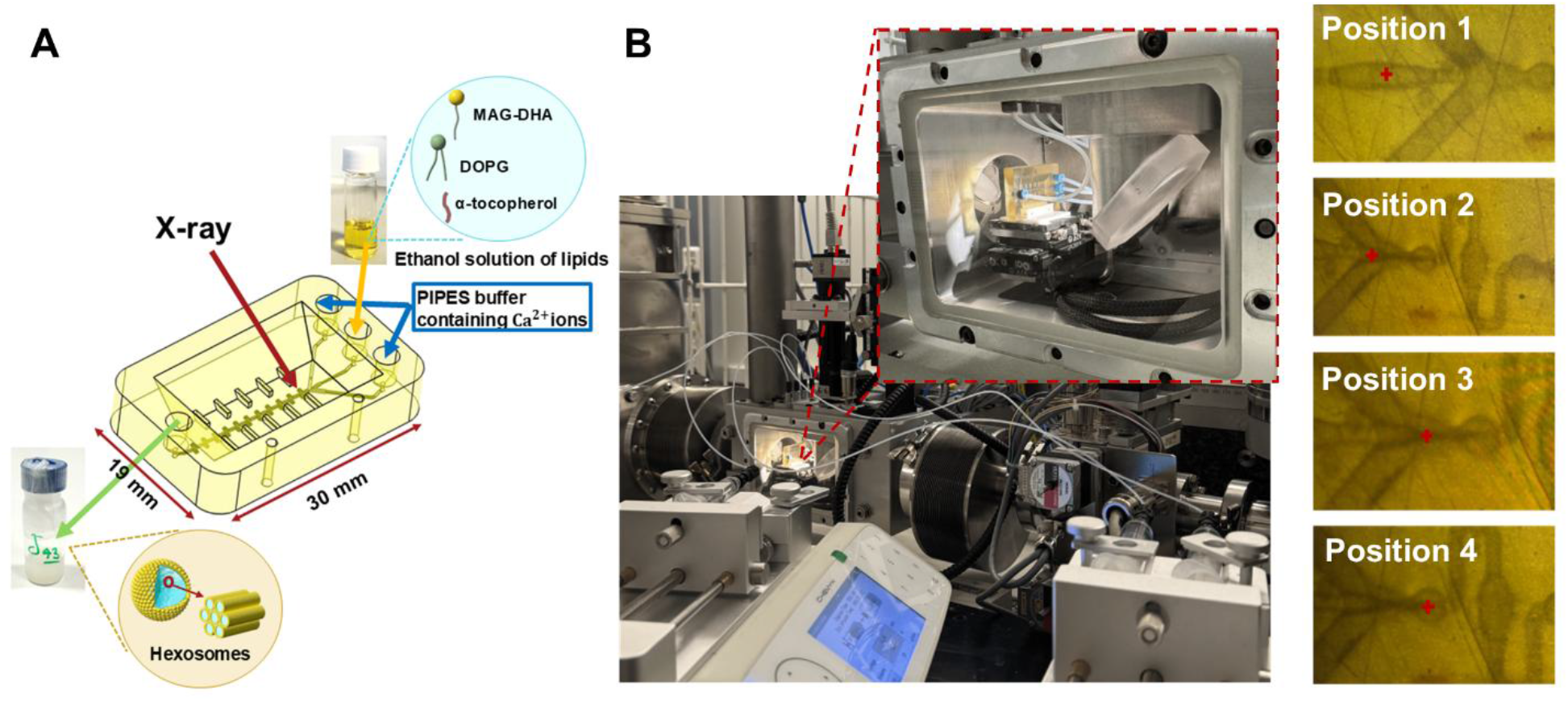
(A) Schematic representation of the 3D-printed hydrodynamic flow-focusing (HFF) chip used for hexosome formation, showing the central ethanolic lipid stream focused by two Ca^2+^-containing PIPES buffer streams and the X-ray probing direction. (B) The HFF chip mounted on a piezo stage inside a vacuum chamber at the P12 beamline (PETRA III, DESY, Hamburg, Germany) for SAXS-on-chip measurements. The enlarged view shows the mounted chip, and the optical micrographs indicate the four selected SAXS measurement positions (1– 4) along the microchannel.

The 3D-printed HFF chip was used for continuous production of nanodispersions (hexosomes) at a fixed total flow rate (TFR) of 200 µLmin^−1^ and the following three different flow rate ratios (FRRs): 10, 20, and 30. It is worth noting that FRR is the flow ratio of the combined two sheath flow streams (side channels) to the central lipid stream. All syringes were mounted on syringe pumps (Fusion 200X, Chemyx Inc., Stafford, TX, USA) and connected to the microfluidic chip using polytetrafluoroethylene (PTFE) tubing (Mikrolab, Aarhus, Denmark) and Mini Luer connectors (Microfluidic ChipShop, Jena, Germany). For subsequent SAXS analysis, the produced nanodispersions were then collected at the outlet into 1.5 mL screw-neck vials (Macherey-Nagel, Düren, Germany). Based on visual inspections, all nanodispersions were found to be colloidally stable for at least one month of storage at 25 °C.

### 2.4. Microfluidic Mixing Dynamics and Time-scale Analysis

Here, the Reynolds number (Re) was estimated to be approximately 15, indicating laminar flow regime in the microchannel, where mixing is governed predominantly by molecular diffusion across fluid interfaces.

For laminar flow, the local residence time *t_res_* at a downstream position *x* relative to the mixing origin (*x* = 0) is defined by the mean flow velocity (*U*) ^13^:

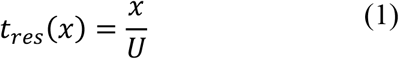

At TFR of 200 µL min^−1^, within the expanded channel section (width *W* = 270 µm; height *H* = 200 µm; cross-sectional area = 0.054 mm^2^), *U* along the central channel was calculated to be 61.7 mm s^−1^. Accordingly, the corresponding *t_res_* values at downstream distances of *x* ≈ 0, 450, and 850 µm were estimated to be approximately 0, 7.3, and 13.8 ms, respectively. These positions correspond to the SAXS measurement points labeled Positions 2-4 in **Figure 2B**.

In hydrodynamic focusing systems, mixing efficiency is primarily governed by the width of the focused solvent stream (*w_f_*), which defines the characteristic diffusion length scale. The focused stream width and the corresponding diffusive mixing time (*τ_mix_*) were estimated as follows^13–15^:

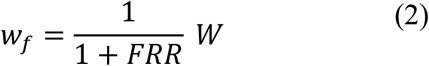

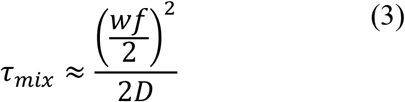

Using the diffusion coefficient of ethanol in water^41^ (*D* = 1.23 × 10^−9^ *m*^2^*s*^−1^) and FRR of 10 the focused stream width was calculated to be *w_f_* ≈24.5 µm, yielding a theoretical *τ_mix_* value of approximately 62.8 ms. This condition corresponds to the one employed in the online SAXS-on-chip experiments.

### 2.5. Small-Angle X-ray Scattering

The online SAXS-on-chip experiments were carried out at room temperature at the P12 synchrotron beamline at EMBL Hamburg (PETRA III, DESY, Hamburg, Germany)^42^, with measurements acquired at different positions along the center channel of the 3D-printed microfluidic chip, as illustrated in **Figure 2B**. The microfluidic chip was mounted inside a vacuum chamber on a piezo motor stage using a custom-designed holder, fabricated *via* fused filament fabrication (FFF) by using a desktop 3D printer (Ender V3, Creality, Shenzhen, China). An on-axis camera was used to precisely align the microfluidic channels with the X-ray beam. The X-ray beam with an energy of 10 keV was collimated to an effective size of approximately 100 × 200 μm^2^ (full width at half maximum, FWHM) at the sample position, delivering a photon flux of 5 × 10^12^ photons s^−1^. Data acquisition was performed using a Pilatus 6 M detector (Dectris Ltd., Baden, Switzerland) at a sample-to-detector distance of 3 m, covering a momentum transfer, *q*, range of 0.02 nm^−1^ < *q* < 4.80 nm^−1^, where *q* = 4π sin θ/λ. 2θ is the scattering angle and λ is the X-ray wavelength of 0.124 nm). Silver behenate (CH_3_-(CH_2_)_20_-COOAg) with a *d*-spacing value of 5.84 nm was used as a standard to calibrate the angular scale of the measured intensity. Each read-out consisted of 10 frames with an exposure time of 95 ms per frame. The 2D scattering images were azimuthally integrated and normalized to the transmitted beam intensity using the automated analysis pipeline SASFLOW, yielding one-dimensional (1D) SAXS profiles, as previously described^43^. Background scattering from PIPES buffer (10 mM, pH 7.0) was measured in the central microfluidic channel prior to sample acquisition. Before conducting the online SAXS-on-chip experiments, background measurements were repeated until a stable and reproducible scattering signal was obtained at all predefined positions along the central channel (**Figure S2**). The slight variations observed in the SAXS patterns during the initial exposures were consistent with beam-induced changes in the resin, potentially involving further polymerization or crosslinking. Once the background scattering stabilized, the corresponding stabilized pattern was used as the background for subsequent measurements acquired at the same position.

In addition to online SAXS-on-chip measurements, the nanodispersions produced under the same experimental conditions were characterized using a laboratory SAXS instrument (BioXolver L, Xenocs, Grenoble, France) equipped with a GeniX 3D Cu Kα X-ray source (λ = 0.134 nm) at 25 °C. The sample-to-detector distance was 0.633 m, covering *q* in the range of 0.20–5.00 nm^−1^. 30 μL of each nanodispersion was manually loaded into a quartz capillary, and five frames were recorded with an exposure time of 60 s per frame. The 2D scattering images were azimuthally integrated by using BioXTAS RAW software^44^ to produce 1D SAXS patterns, which were then averaged background-subtracted using PIPES buffer.

To characterize the internal nanostructure of the produced nanodispersions, the *q*_ℎ*k*_ positions of the detected Bragg reflections were determined by Lorentzian fitting, and the corresponding *d*_ℎ*k*_ values were calculated using *d*_ℎ*k*_ = 2*π*/*q*_ℎ*k*_. An internal inverse hexagonal (H_2_) was identified from the characteristic reflection ratios 1: √3: √4, …^45–48^. The corresponding lattice parameter, *a*, was derived by the hexagonal scattering law, 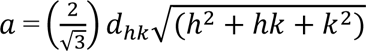 where (h,k) are the Miller indices of the corresponding reflections^45,46,48^. All reflection analysis and structural parameter calculations were performed custom Python scripts.

### 2.6. Cryo-Transmission Electron Microscopy

Cryo-transmission electron microscopy (Cryo-TEM) measurements were performed on selected nanodispersions as previously described^49,50^. Briefly, 3 μL of each sample was deposited on a hydrophilized 300-mesh lacey carbon copper grid (Ted Pella Inc., Redding, CA, USA). The nanodispersion was then blotted with filter paper using a Vitrobot (FEI, Eindhoven, The Netherlands) for 6 s at 22 °C and 100% humidity, with a blotting force of 6. Immediately afterward, the nanodispersion was vitrified by plunging into liquid ethane cooled by liquid nitrogen. Imaging was performed using a Tecnai G2 20 TWIN transmission electron microscope (FEI, Eindhoven, The Netherlands) operated at 200 kV under low-dose conditions.

### 2.7. Dynamic Light Scattering Measurements

Dynamic light scattering (DLS) measurements were conducted to determine the mean nanoparticle size and the polydispersity index (PDI), which reflects the nanoparticle size distribution, by using a Zetasizer Nano ZS system (Malvern Instruments Ltd., Worcestershire, UK) equipped with a 633 nm laser and 173° backscatter detection optics. Prior to measurements, each nanodispersion was diluted 100-fold in PIPES buffer (10 mM, pH 7.0) alone or PIPES buffer (10 mM, pH 7.0) containing 5 mM Ca^2+^ ions. The diluted samples were subsequently filtered through hydrophilic nylon syringe filters (0.22 µm, Frisenette, Knebel, Denmark). All measurements were conducted in triplicate at 25 °C following 2 min of equilibration. The dispersant viscosity and refractive index were set to the value for water at 25 °C. The data was processed using Malvern Zetasizer software v7.12 (Malvern Instruments Ltd., Worcestershire, UK).

### 2.8. Molecular Dynamics Simulations

#### 2.8.1. Simulation Methodology

Coarse-grained molecular dynamics (CG-MD) simulations were performed using the Martini 3 force field^51^ within GROMACS (2023.5)^52^. Multilamellar lipid systems (15 × 15 × 6 nm³) were mimicked by the stacking of two identical lipid bilayers. The bilayers were generated using the INSANE tool ^53^ and subsequently replicated along the z-axis using the *gmx genconf* function^52^ to obtain the final systems. Each system was subsequently solved with approximately 3-5 Martini 3 water beads per lipid, corresponding to 12-20 water molecules per lipid, as each Martini water bead represents four atomistic water molecules. Simulations were performed for MAG-DHA/DOPG/α-tocopherol (48:40:12, w/w/w), MAG-DHA/DOPG (37:63, w/w), and MAG-DHA/DOPG/α-tocopherol (34:57:9, w/w/w) systems in the presence and absence of α-tocopherol and Ca^2+^ ions, using a Ca^2+^/DOPG molar ratio of 0.5 when Ca^2+^ ions were included. The MAG-DHA/DOPG (34:57:9) system was simulated both with and without Ca^2+^ ions to assess the role of Ca^2+^–DOPG interactions in modulating the structural features. Lipid parameters for MAG-DHA and DOPG were obtained from the recently published Martini 3 lipidome library^54^.

Prior to simulation, the systems underwent energy minimization for 500 steps using the steepest descent algorithm to remove any unfavorable clashes or overlaps between beads. This was followed by a 10 ns relaxation phase to allow the molecules to settle into a stable arrangement. Finally, production simulations were performed for 2 µs in triplicate. During both equilibration and production, the temperature was kept constant at 25 °C. Temperature coupling was achieved using the Berendsen thermostat^55^ during equilibration and the velocity-rescale thermostat^56^ during production, with a coupling constant of 1 ps. Lipids and the water– NaCl solution were coupled to separate temperature baths.

The systems were maintained at constant pressure using anisotropic pressure coupling during both the relaxation and production runs, using the Berendsen^55^ and Parrinello-Rahman^57^, respectively. Pressure was coupled with a coupling constant of 12 ps and compressibility at 3×10^−4^ bar^−1^.

#### 2.8.2 Parametrization of the α-tocopherol Coarse-Grain Model

Since prior to this work no existing model for α-tocopherol was available in the Martini 3 lipidome library^54^, new parameters were generated in this study. Using the software LigPargen with the OPLS-AA/1.14*CM1A(-LBCC) force field^58^, atomistic parameters for α-tocopherol were first generated. Then, the obtained fragment was simulated for 1 µs in 150 mM NaCl solution. This fragment was energy-minimized using steepest descent algorithm with 5000 steps, followed by a relaxation of 250 ps using a 1 fs timestep, and a production run of 1 µs using a 2 fs time step. The temperature and pressure were held constant at 25 °C and 1 bar, respectively, using Berendsen^55^ thermostat and barostat for relaxation and V-rescale thermostat^56^ and Parrinello-Rahman barostat^57^for the production run.

Atomistic simulations were mapped into coarse-grained (CG) resolution using MDAnalysis^59,60^. The mapping for α-tocopherol is shown in **Figure S3A**. The aromatic core was constructed as a rigid body using three constraints, with additional interaction sites defined as type 3 virtual sites as previously described for other Martini 3 molecules^53^. The tail mapping and bead choices were based on Koukos et al.^61^. To ensure an accurate description of the molecular shape and volume, the Connolly surface of the CG model (**Figures S3B**) and the solvent accessible surface area (**Figure S3C**) were validated by comparing with all-atom MD results for the same molecules in solution. α-tocopherol was then simulated in CG detail for 1 µs in 150 mM NaCl solution. The bonded distributions were fitted onto the corresponding atomistic distributions (**Figure S4**). Due to the hydrophobicity of α-tocopherol, a simulation in chloroform was performed as well to extract the behaviour of the molecule in a hydrophobic environment, similar to that of a membrane. For both the CG simulations in the aqueous NaCl solution and those in chloroform, the system was minimized for 500 steps using the steepest descent algorithm. This was followed by a 10 ns relaxation by using a 10 fs timestep, and finally a 1 µs production run with a 20 fs time step. The temperature and pressure were kept constant during both the relaxation and production runs at 25 °C and 1 bar, respectively. The pressure coupling was isotropic and achieved through use of Berendsen^55^ and Parinello-Rahman barostats^57^, respectively, during the relaxation and production runs, with the compressibility set to 3×10^−4^ bar^−1^. Berendsen^55^ and Velocity-rescale thermostat^56^ were used, respectively, for relaxation and production runs with a coupling constant of 1.0 ps for maintaining the temperature at 25 °C.

#### 2.8.3 Analysis of Molecular Dynamics Simulations

To evaluate the time evolution of α-tocopherol preference for DOPG or MAG-DHA, intermolecular contacts were calculated using a distance-based criterion. A contact was defined as any instance in which a bead of α-tocopherol was within 0.47 nm of any bead of the lipid (DOPG or MAG-DHA). For each trajectory frame, the number of α-tocopherol–DOPG and α-tocopherol–MAG-DHA contacts was computed.

The number of contacts was normalized by the total number of beads in each lipid species. The enrichment factor was defined as the ratio of the normalized contacts for every frame:

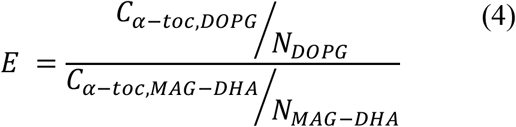

where (E) is the enrichment ratio, (C) is the number of contacts, and (N) is the number of beads for the indicated lipid species.

Ca^2+^–lipid interactions were quantified using a distance-based criterion. A Ca^2+^ ion was considered to interact with a lipid, either DOPG or MAG-DHA, when it was located within 0.47 nm of the lipid headgroup bead. For each trajectory frame, pairwise distances between Ca^2+^ ions and lipid headgroup beads were computed. The mean number of coordinating lipids per Ca^2+^ ion was separately calculated for DOPG and MAG-DHA and monitored as a function of time for each replicate.

Both MD calculations were performed using were performed using in-house Python scripts leveraging MDAnalysis^59,60^ and numpy libraries^62^.

### 2.9. SAXS Calculation from Coarse-Grained Martini Structures

SAXS profiles were computed from individual coarse-grained configurations using a direct reciprocal-space formulation and were subsequently ensemble-averaged over the sampled configurations. The input structures were provided as Protein Data Bank (PDB) files and were first checked to ensure that the atom name field encoded the bead type instead of the regular bead name. The change was required for the heuristic evaluation of the electron count in the scattering calculation. Throughout the calculation, the simulation cell was treated as periodic.

Each bead was represented as a homogeneous sphere, with bead-type-dependent effective electron counts assigned to define the zero-angle scattering intensity. Thus, the coarse-grained beads were treated as finite scatterers, analogous to conventional SAXS, with the scattering contribution of each site modulated by its form factor rather than approximated as that of a point scatterer. An effective bead radius was therefore assigned to each bead type, allowing the finite spatial extent of the coarse-grained sites to be retained in the scattering model. Under this representation, the *q*-dependent form factor of bead *i* was expressed as:

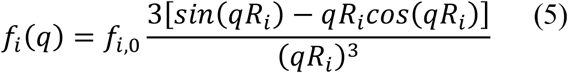

where *f_i,_*_0_ is the zero-angle scattering strength and *R_i_* is the effective bead radius. This analytical sphere form factor was used for all beads in the system.

Because the structures were interpreted as periodic bulk systems, the scattering amplitudes were evaluated on the reciprocal lattice of the simulation box with dimensions *L_x_*, *L_y_*, and *L_z_*.

The reciprocal-space vectors were defined as: 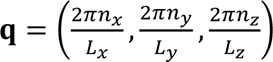, where *n_x_*, *n_y_*, and *n_z_* are integers. Prior to the Fourier evaluation, all bead coordinates were wrapped into the primary simulation cell. For each reciprocal-space vector **q**, the total scattering amplitude was then computed as:

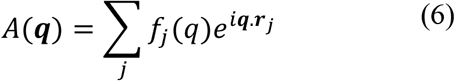

with **r***_j_* denoting the position of bead *j*. The corresponding intensity follows directly from the squared modulus of this amplitude, *I*(***q***) = | < *A*(***q***)^2^ > |.

To obtain the one-dimensional SAXS profile, the discrete reciprocal-lattice vectors were grouped into radial shells according to the magnitude of the scattering vector, *q* = |***q***|, using a uniform shell thickness of Δ*q*. The center of each shell was taken as the corresponding radial scattering value, (*q_k_*). The scattering intensity at *q_k_* was calculated as the average of the intensities of all reciprocal-lattice vectors whose magnitudes fell within the corresponding shell. This scheme leads to a low sampling of reciprocal space, for low *q*. To remedy this, the one-dimensional profile was smoothed along the *q*-axis using a normalized Gaussian kernel with a width of *σ_q_* = 0.75Δ*q*. This smoothing was applied only as a post-processing step and did not alter the underlying reciprocal-lattice evaluation. The lower bound of the meaningful accessible scattering range is 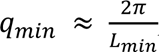, below which correspond to wavelengths larger than the shortest dimension of the periodic cell (*L_min_*) and are therefore particularly sensitive to finite-size effects.

The calculation was performed using an in-house script called martini_saxs command-line tool which is available at https://github.com/fabianschuhmann/martini_saxs. Here, the scattering range up to *q*_max_ = 3.5 nm^−1^ was used. The 1D profile was obtained by radial averaging onto *N_q_* = 80 uniformly spaced target *q*-values over the selected *q*-range, where *N_q_* denotes the number of points in the output *q*-grid. The corresponding grid spacing was denoted by Δ*q*.

## 3. RESULTS AND DISCUSSION

### 3.1. Experimental Validation and Surface Characterization of the 3D-Printed Microfluidic Chip

Accurate microfluidic channel dimensions with consistency between fabricated microfluidic structures and their design specifications are essential. Accurate microfluidic channel dimensions are vital for ensuring consistency between fabricated microfluidic structures and their design specifications. Any alterations in these dimensions may affect the fluid flow parameters and compromise reproducibility of experiments^63,64^. In this study, the dimensional accuracy of the fabricated microfluidic channels was validated in three complementary approaches. First, fluorescence microscopy (Nikon Eclipse LV150N, Nikon Corporation, Tokyo, Japan) was used to monitor fluorescently labeled nanoparticles in transit through the microfluidic chip. As shown in **Figure 3A**, the micrographs exhibited uniform fluorescence intensity throughout the serpentine paths and junctions, confirming continuous and unobstructed flow through the printed microchannels. Measurements at multiple locations along the channels showed widths of 97–107 µm and 269–275 µm for nominal dimensions of 100 and 270 µm, respectively. The close correspondence between the measured and designed values demonstrates good lateral dimensional fidelity and consistency of the 3D-printing process. Second, scanning electron microscopy (SEM, Zeiss Supra VP 40, Oberkochen, Germany) was employed to examine the structural details of a half-printed microchannel after sputter-coating with a 10 nm gold layer (Cressington 208HR, Watford, UK). The representative SEM image (**Figure 3B**), together with the high-magnification inset, clearly resolved the individual printed layers, demonstrating high 3D printing fidelity that allows for a precise estimation of the printed layer thickness. Approximately 10 distinct layers were clearly distinguishable across a measured channel height of ∼45 µm at the mixing junction. After accounting for the 30° imaging tilt, the corresponding channel height was estimated to be approximately 52 µm, yielding an average layer thickness of approximately 5.2 µm. This value closely agrees with the intended layer thickness of 5 µm in the channel region, supporting good dimensional fidelity in the vertical direction. Finally, a 3D surface profile was acquired using confocal laser scanning microscopy (Keyence VK-X3000, Osaka, Japan). As illustrated in the volumetric reconstructions in **Figure 3C**, the topographic maps provided a height-coded visualization of the channel architecture. The distinct color gradient between the upper surface and the bottom wall of the microfluidic channel, combined with the isometric 3D view (bottom panel), confirmed the uniform channel depth and well-defined wall verticality across the scanned area (approximately 2765 µm ×2073 µm).

**Figure 3.**
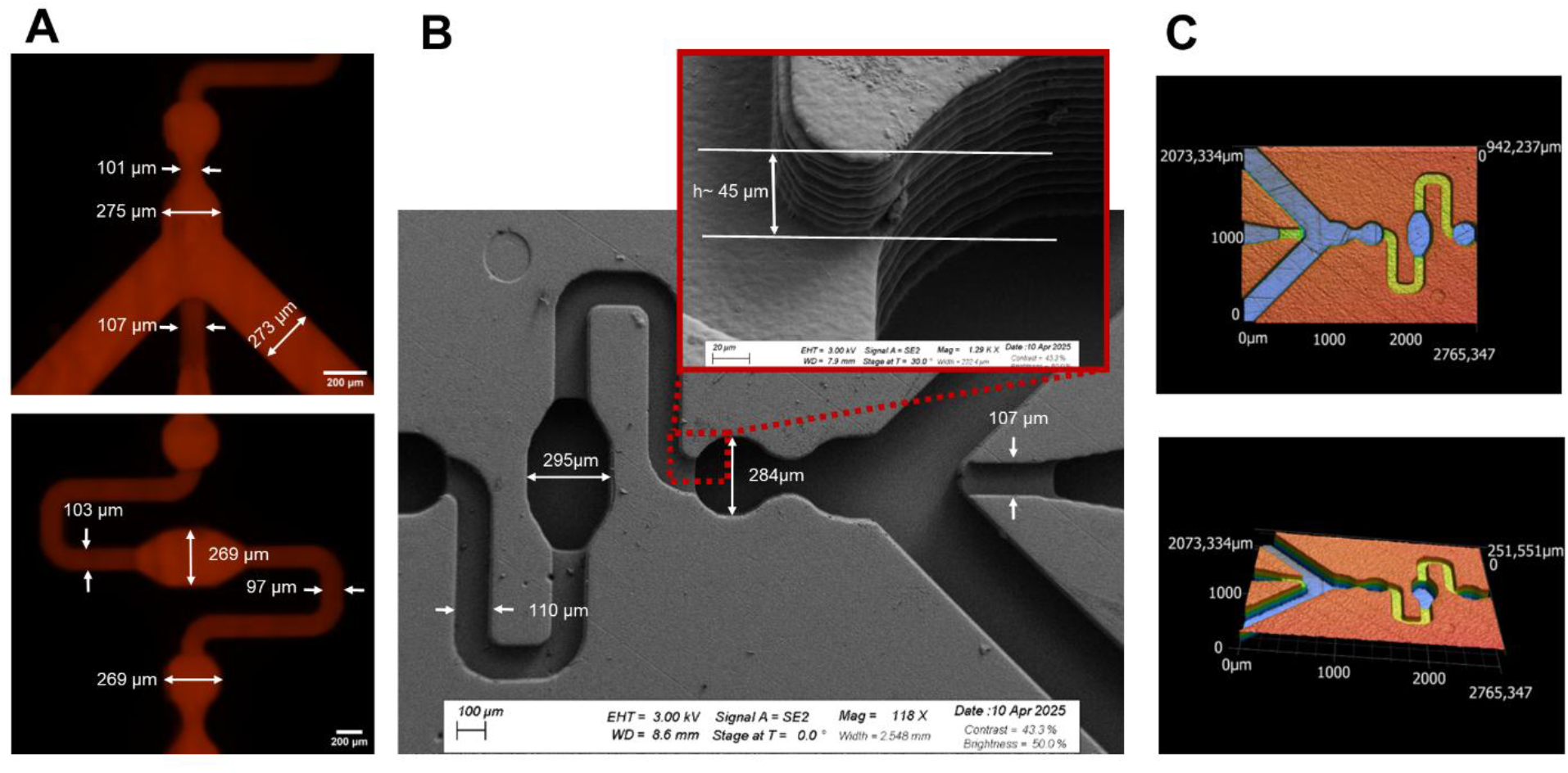
Dimensional validation of the 3D-printed microfluidic channel. (A) Fluorescence microscopy image of the channel filled with fluorescent nanoparticles, showing continuous filling throughout the serpentine paths and junctions. Measurements obtained at multiple locations yielded channel widths of 97–107 µm and 269–275 µm for nominal dimensions of 100 and 270 µm, respectively. (B) Scanning electron microscopy (SEM) image of a half-printed channel after deposition of a 10 nm gold layer. The high-magnification image resolves approximately 10 individual printed layers across a measured channel height of ∼45 µm at the mixing junction. After correction for the 30° imaging tilt, the estimated channel height was ∼52 µm, corresponding to an average layer thickness of ∼5.2 µm, close to the nominal 5 µm layer thickness. (C) 3D surface profile of the half-printed channel obtained by confocal laser scanning microscopy (red = top surface, blue = channel bottom, yellow = channel bottom of serpentine region). The distinct height contrast between the upper surface and the channel bottom, together with the isometric 3D view (bottom panel), confirmed a uniform channel depth and well-defined, nearly vertical sidewalls across the scanned area (approximately 2765 µm ×2073 µm).

In addition to the microfluidic channel dimensions, the surface properties of the HTL resin play a crucial role in modulating the microfluidic performance as they determine the microchannel wettability and fluid dynamics within the channels^65–68^. It was, therefore, essential to assess the surface roughness and wettability of the bulk HTL resin used in the chip fabrication process. The arithmetic average roughness (*R_a_*) was measured on a representative flat-printed surface of the resin by using laser confocal scanning microscopy (Keyence VK-X3000, Osaka, Japan). The analysis yielded a mean *R_a_* value of 42 ± 1 nm. This low-profile roughness level indicated that the printing process produces exceptionally smooth surfaces. This is essential for minimizing hydrodynamic instabilities and promoting a stable predictable laminar flow within the 3D-printed microfluidic chip. Furthermore, static contact angle measurements were conducted using a Drop Shape Analyzer (KRÜSS GmbH, Hamburg, Germany) to evaluate the surface wettability of the HTL resin. The obtained mean contact angle of 63.91 ± 0.61° confirmed a moderate hydrophilicity of the HTL resin surface. Such surface characteristics are well-suited for aqueous-phase microfluidic platforms designed for continuous production of colloidal nanoobjects^69^. They enable reliable filling of the channel network with aqueous media and provide a favorable environment for achieving consistent fluid–wall interactions without inducing excessive solute adhesion^70^. For continuous production of lipid nanoparticles, these surface properties are particularly important for preventing lipid adherence to the channel walls^26^. In addition to a successful collection of high quality SAXS data and their analysis, they facilitated cleaning and handling of the 3D-printed microfluidic chips between the subsequent SAXS experiments.

### 3.2. SAXS-on-Chip Study of Hexosome Formation Dynamics

To gain insight into the dynamic self-assembly behavior during the continuous production of hexosomes, online SAXS-on-chip measurements were performed using the 3D-printed microfluidic chip during controlled mixing of an ethanolic solution containing the ternary MAG-DHA/DOPG/α-tocopherol lipid mixture with PIPES buffer (10 mM, pH 7.0) containing 5 mM Ca^2+^ ions. The molecular structures of the lipids MAG-DHA and DOPG, along with α-tocopherol, are presented in **Figure 4**. Two lipid compositions, MAG-DHA/DOPG/α-tocopherol 48:40:12 and 56:30:14 (w/w/w), were studied under hydrodynamic flow-focusing conditions (FRR = 10; TFR = 200 μL min^−1^).

**Figure 4.**
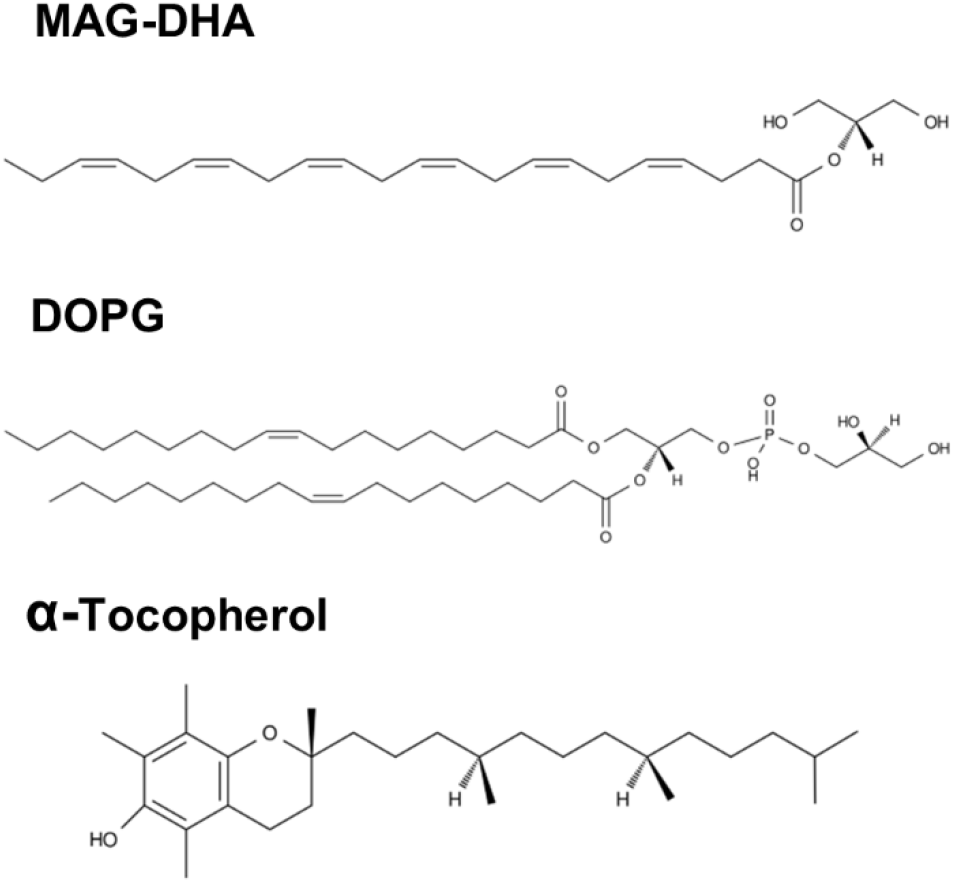
Molecular structures of docosahexaenoic acid monoacylglycerol (MAG-DHA), 1,2-dioleoyl-sn-glycero-3-phosphoglycerol, (DOPG) and α-tocopherol.

At position 1 on the microfluidic chip (**Figure 2B**), before contact with the Ca^2+^-containing buffer and prior to substantial solvent exchange, the online SAXS patterns obtained for both MAG-DHA/DOPG/α-tocopherol compositions showed diffuse scattering, consistent with the presence of micellar solutions (**Figure 5A,B**). This is consistent with studies reporting that biologically relevant amphiphilic lipids dissolved in ethanol can form normal micellar solutions at relatively high ethanol concentrations^12,71,72^. Therefore, these measurements reflect the initial ethanol-rich state of the lipid solutions prior to buffer-induced solvent exchange. After exposure of the ethanolic lipid solution to the Ca^2+^-containing buffer, SAXS patterns collected at different positions along the microfluidic channel were used to follow the evolution of the internal nanostructure. For the MAG-DHA/DOPG/α-tocopherol 48:40:12 (w/w/w) composition, a very fast colloidal transformation was already evident at position 2 (x ≈ 0), corresponding to the earliest probed point after buffer exposure. The initially micellar ethanol-rich solution was already transformed into nanoparticles with a biphasic internal feature comprising two coexisting inverse hexagonal (H_2_) phases (**Figure 5A**). These two coexisting H_2_ phases were identified with lattice parameters of 5.95 and 6.54 nm, corresponding to reflections at *q* values of approximately 1.22, 2.11, and 2.44 nm^−1^ and 1.11 and 1.92 nm^−1^, respectively. This near-immediate micelle-to-hexosome transformation highlights the rapidity of solvent-exchange-driven lipid self-assembly in the microfluidic channel. Here, the early and rapid colloidal closure into nanoparticles with a biphasic internal architecture comprising two coexisting H_2_ phases likely reflects transient compositional heterogeneity arising from rapid ethanol diffusion into the excess aqueous buffer and concomitant redistribution of ethanol, water, and lipid components at the water–ethanol interfacial region. Such nonequilibrium interfacial gradients^71^ may generate local variations in spontaneous curvature and thereby facilitate the early formation of the two coexisting H_2_ phase within the first milliseconds of the self-assembly process.

**Figure 5.**
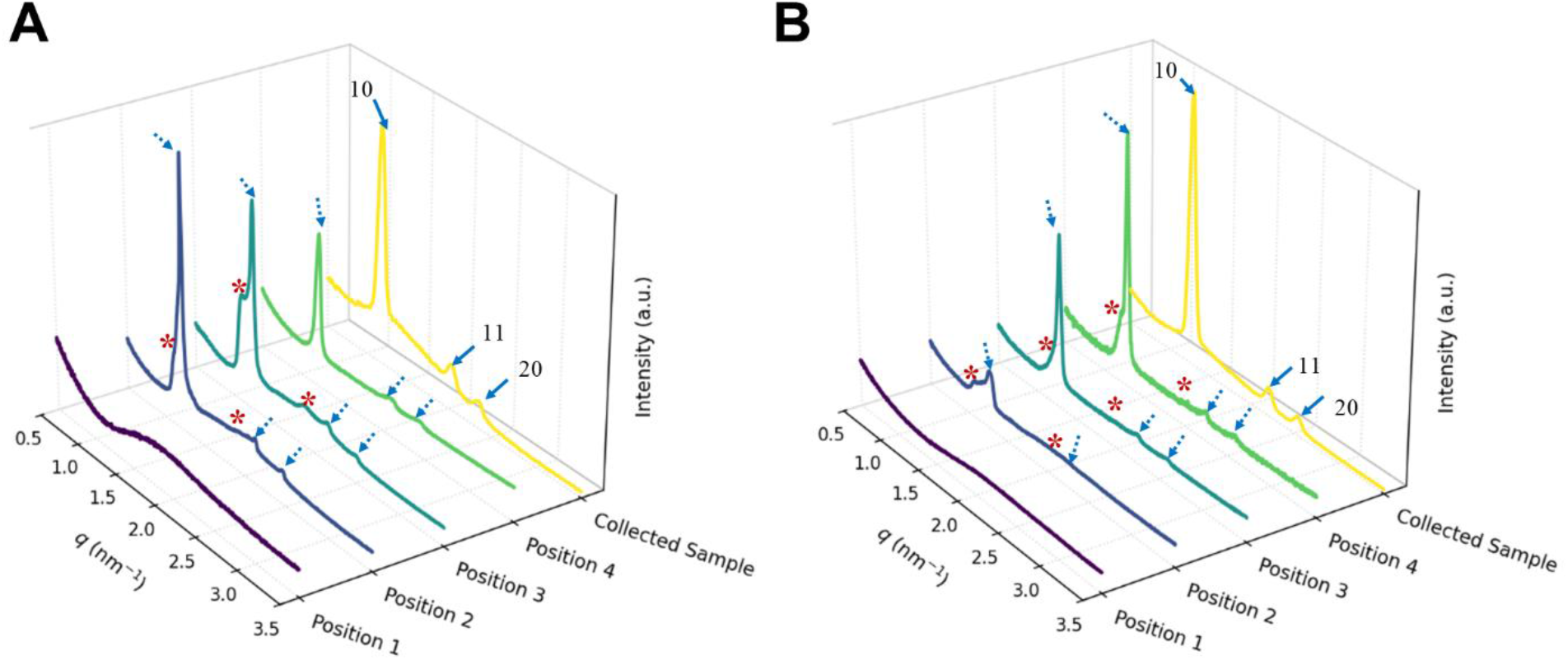
SAXS patterns recorded along the microchannel at FRR 10 and a constant TFR of 200 µL min^−1^ for MAG-DHA/DOPG/α-tocopherol at weight ratios of (A) 48:40:12 (w/w/w) and (B) 56:30:14 (w/w/w). The synchrotron scattering intensities from online SAXS-on-chip measurements were normalized to the scattering intensity of the collected nanodispersions (yellow SAXS atterns) obtained using a laboratory SAXS instrument to enable direct comparison. Miller indices are indicated by blue arrows. Dashed blue arrows and red stars indicate two coexisting H2 phases during online SAXS-on-chip measurements.

The internal biphasic H_2_ architecture persisted at position 3 (*t*_res_ ≈ 7 ms), where the two coexisting H_2_ phases were retained with lattice parameters of 5.85 and 6.57 nm (**Figure 5A**). Compared with position 2, a slight decrease in the lattice parameter of the H_2_ phase, initially observed at approximately 5.95 nm, was noted, while the coexisting H_2_ phase with a lattice parameter of around 6.5 nm remained essentially unchanged. By position 4 (*t_res_* ≈ 14 ms), the biphasic feature progressively evolved toward a neat internal H_2_ phase with a lattice parameter of 6.15 nm, with corresponding reflections observed at *q* values of approximately 1.17, 2.04, and 2.37 nm^−1^. This structural evolution suggests that ethanol diffusion into the excess aqueous buffer, together with progressive redistribution of ethanol, water, and lipid components, ultimately leads to the formation of hexosomes with a neat internal H_2_ phase, likely reflecting a more uniform distribution of lipids and residual ethanol within the self-assembled interiors of the nanoparticles. Thus, the early appearance of well-defined H_2_ Bragg reflections, at times substantially shorter than the estimated diffusive mixing time of approximately 63 ms (Section 2.4), further suggests that hexosome formation is governed predominantly by interfacially driven nonequilibrium processes rather than by complete solvent equilibration. A similar early micelle-to-hexosome colloidal transformation was observed for the MAG-DHA/DOPG/α-tocopherol 56:30:14 w/w/w composition (**Figure 5B**). At position 2, two coexisting H_2_ phases were again identified, with lattice parameters of 5.72 and 6.77 nm. In contrast to the 48:40:12 w/w/w composition, this biphasic H_2_ feature persisted through positions 3 and 4, with only moderate changes in the corresponding lattice parameters of both phases. The H_2_ phase initially observed at approximately 5.72 nm evolved within the range of approximately 5.61-5.64 nm, whereas the lattice parameter of the coexisting H_2_ phase initially observed at approximately 6.77 nm shifted toward approximately 5.98-6.10 nm. The corresponding *q*-values and lattice parameters obtained at each channel position are presented in **Table S1.**

Together, the two online SAXS-on-chip experiments (**Figure 5**) show that the structural evolution of the hexosomes is composition-dependent and is not governed solely by Ca^2+^– DOPG interactions. Ca^2+^ association with anionic phospholipids such as DOPG has been proposed to proceed through two stages: first, enrichment of Ca^2+^ at the lipid–water interface, and second, formation of coordination complexes with the anionic DOPG headgroups^73,74^. Such Ca^2+^–DOPG interactions are expected to promote hexosome formation by screening the negative charge of the DOPG headgroups, reducing their effective headgroup area through interfacial dehydration, and promoting tighter packing within the interfacial film, as discussed in previous studies^26,73–75^. These Ca^2+^-mediated interfacial dehydration and condensation effects most likely facilitate the transformation from the initially micellar ethanol-rich state to nonlamellar colloidal nanoparticles. However, the formation of hexosomes also strongly depends on lipid composition. In both experiments, MAG-DHA contributes to the lamellar-to-nonlamellar phase transition because of its intrinsic propensity to form an H_2_ phase in excess water, whereas DOPG tends to form a lamellar phase^76,77^. Thus, the DOPG/MAG-DHA ratio, the corresponding DOPG/Ca^2+^ molar ratio, lipid–lipid and lipid–water interactions, and solvent-exchange-driven interfacial gradients collectively modulate the structural features of the produced nano-self-assemblies.

To further assess the structural evolution associated with the microfluidic production process, collected nanodispersions corresponding to the two lipid compositions and prepared under the same flow conditions as the online SAXS-on-chip experiments were characterized *ex situ* by SAXS (yellow patterns in **Figure 5A,B**). For both lipid compositions, comparison of the *ex situ* SAXS patterns of the final collected nanodispersions with those obtained on-chip at position 4 under nonequilibrium conditions revealed a slight increase in the lattice parameter of the internal H_2_ phase of the produced hexosomes. For the MAG-DHA/DOPG/α-tocopherol compositions of 48:40:12 and 56:30:14 (w/w/w), the lattice parameter of the neat internal H_2_ phase increased from approximately 6.15 to 6.22 nm and from approximately 5.64 to 5.73 nm, respectively (**Table S1**). As discussed above, this small structural difference indicates that the internal H_2_ phase progressively evolves along the microfluidic channel as ethanol diffuses out of the lipid-rich stream and water penetrates the forming colloidal nanoparticles. Thus, the on-chip structure detected at position 4 represents an advanced nonequilibrium state that has begun to approach the neat internal H_2_ phase of the final collected hexosomes.

The SAXS patterns of the collected nanodispersions displayed a more pronounced diffuse scattering in the low-q regime than the corresponding SAXS-on-chip patterns recorded at positions 2-4 (yellow patterns in **Figure 5A,B**). This suggests that the collected nanodispersions contained a higher fraction of weakly ordered coexisting structures. This contribution most likely arises from DOPG-rich vesicles formed when part of the DOPG partitions into the buffer upon direct exposure of the ethanolic ternary lipid solution. As solvent exchange proceeds along the center microfluidic channel, the fraction of these coexisting vesicles appears to increase, leading to a stronger vesicular contribution in the final collected nanodispersions than during the SAXS-on-chip measurements. Such structural heterogeneity, arising from the coexistence of hexosomes and vesicular structures, is commonly observed in nonlamellar liquid crystalline nanodispersions, particularly in phospholipid-containing nanodispersions where the vesicular contribution is often more pronounced ^76,78^.

To complement the online SAXS-on-chip investigations, offline SAXS experiments were performed to further elucidate the effect of Ca^2+^ concentration on the Ca^2+^-triggered vesicle-to-hexosome colloidal transformation. MAG-DHA/DOPG/α-tocopherol nanodispersions with lipid compositions of 48:40:12 and 56:30:14 were prepared in PIPES buffer (10 mM, pH 7.0) containing 0, 1, 3, or 5 mM Ca²⁺ at an FRR of 10 and a TFR of 200 µLmin^−1^. Additional nanodispersions were prepared at 5 mM Ca^2+^ using FRRs of 20 and 30 to evaluate the effect of increased aqueous dilution on the lipid-to-Ca^2+^ molar ratios. The final Ca^2+^ concentrations after microfluidic mixing and the corresponding DOPG/Ca^2+^ and (DOPG+MAG-DHA)/ Ca^2+^ molar ratios are summarized in **Table 1**.

**Table 1.** Final Ca^2+^ concentrations and molar ratios of DOPG/Ca^2+^ and (DOPG+MAG-DHA)/Ca^2+^ for MAG-DHA/DOPG/α-tocopherol at 48:40:12 and 56:30:14 (w/w/w) after microfluidic mixing with calcium-containing buffer at different FRRs and constant TFR 200 µLmin^−1^.

| Lipid System (wt%) | FRR | $\text{Ca}^{2+}$ in buffer (mM) | Final $\text{Ca}^{2+}$ after mixing (mM) | Total lipid (% w/w) | DOPG/$\text{Ca}^{2+}$ | (DOPG + MAG-DHA)/ $\text{Ca}^{2+}$ |
| --- | --- | --- | --- | --- | --- | --- |
| 48:40:12 | 10 | 0 | 0 | 2.79 | - | - |
|  | 10 | 1 | 0.91 | 2.79 | 15.2 | 51.4 |
|  | 10 | 3 | 2.73 | 2.79 | 5.1 | 17.1 |
|  | 10 | 5 | 4.55 | 2.79 | 3.0 | 10.3 |
|  | 20 | 5 | 4.76 | 1.45 | 1.5 | 5.1 |
|  | 30 | 5 | 4.84 | 0.98 | 1.0 | 3.4 |
| 56:30:14 | 10 | 0 | 0 | 2.88 | - | - |
|  | 10 | 1 | 0.91 | 2.88 | 11.8 | 55.3 |
|  | 10 | 3 | 2.73 | 2.88 | 3.9 | 18.4 |
|  | 10 | 5 | 4.55 | 2.88 | 2.4 | 11.1 |
|  | 20 | 5 | 4.76 | 1.50 | 1.2 | 5.5 |
|  | 30 | 5 | 4.84 | 0.97 | 0.8 | 3.7 |

In the absence of Ca^2+^ ions, the SAXS patterns of both MAG-DHA/DOPG/α-tocopherol nanodispersions were mainly dominated by diffuse scattering (**Figure 6A,B**). This scattering profile is characteristic of weakly correlated bilayers, such as unilamellar (ULVs) and oligolamellar (OLVs) vesicles. A similar SAXS pattern was detected after microfluidic mixing with buffer containing 1 mM Ca^2+^ ions. At this condition, the final Ca^2+^ concentration was 0.91 mM, and the DOPG/Ca^2+^ molar ratios were 15.2 and 11.8 for the MAG-DHA/DOPG/α-tocopherol 48:40:12 and 56:30:14 compositions, respectively. These relatively high ratios indicate that the Ca^2+^ concentration was insufficient to effectively screen the electrostatic repulsion between neighbouring DOPG molecules and induce a lamellar-to-nonlamellar phase transition.

**Figure 6.**
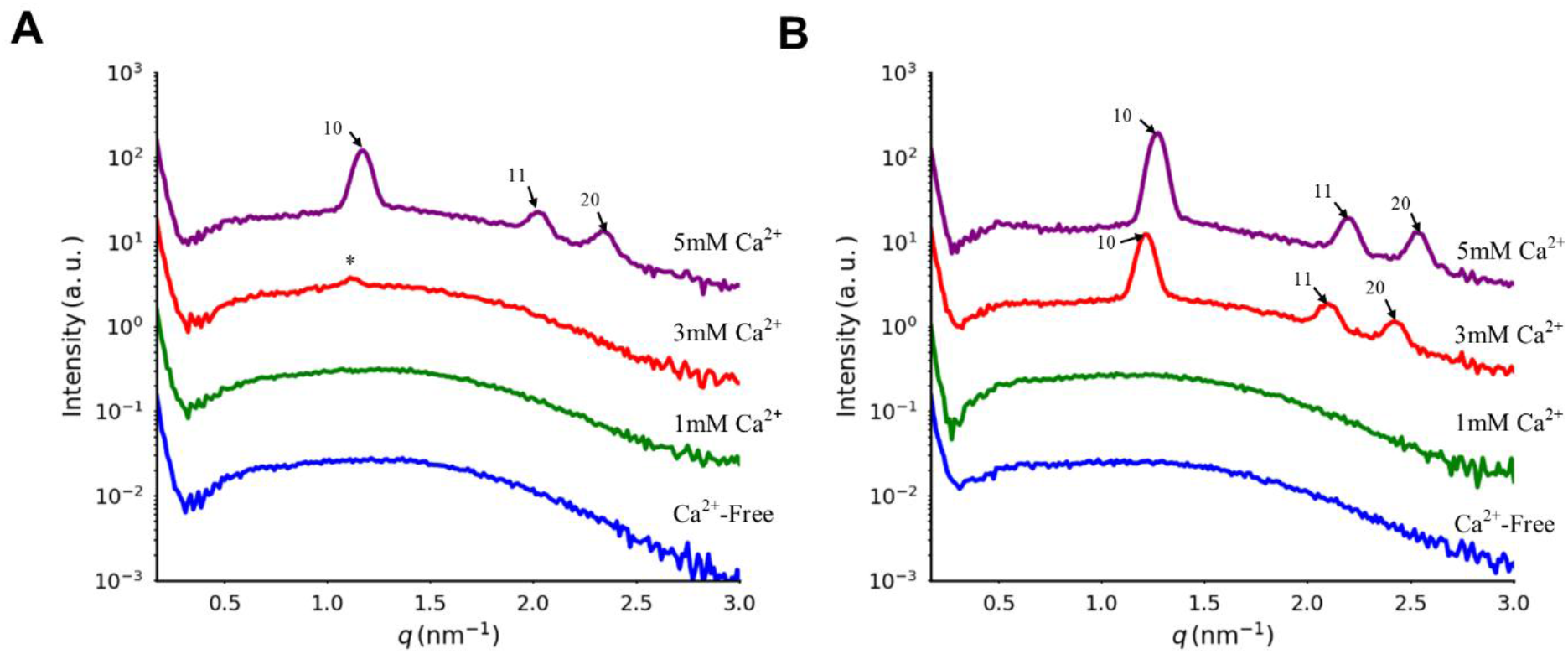
Ca^2+^-induced SAXS patterns of MAG-DHA/DOPG/α-tocopherol nanodispersions at varying Ca^2+^ concentrations of 0,1,3 and 5 mM. Nanodispersions contained MAG-DHA/DOPG/α-tocopherol at weight ratios of (A) 48:40:12 and (B) 56:30:14 (w/w/w) and were prepared by microfluidic mixing at an FRR of 10 and a TFR of 200 µL min^−1^. Miller indices are marked with black arrows. The black star in (panel A) most likely represents the onset of H_2_ phase.

Increasing the Ca^2+^ concentration to 3 mM affected the structural features of the nanodispersions in a composition-dependent manner. For the DOPG-richer composition, MAG-DHA/DOPG/α-tocopherol 48:40:12 (w/w/w), prepared at FRR 10, the SAXS pattern remained largely dominated by diffuse scattering, indicating that vesicular structures were still the dominant population. A weak peak at *q* ≈ 1.2 nm^−1^, marked with an asterisk in **Figure 6A**, was also detected. Because this weak peak appeared at nearly the same *q*-value as the first characteristic peak of the H_2_ phase observed at 5 mM Ca^2+^, as described below, it most likely indicates the presence of traces of the H_2_ phase. Under this condition, the final Ca^2+^concentration in the produced nanodispersion was 2.73 mM and the DOPG/Ca^2+^ molar ratio was 5.1. Upon increasing the Ca^2+^ concentration to 5 mM, the final Ca^2+^ concentration increased to 4.55 mM, and the DOPG/Ca^2+^ molar ratio decreased to 3.0, corresponding to approximately one Ca^2+^ ion per three DOPG molecules. Under this condition, more pronounced Bragg reflections indexed to the H_2_ phase [(10), (11), and (20), marked with black arrows in **Figure 6A**] were detected at *q* values of approximately 1.17, 2.02, and 2.34 nm^−1^, confirming the Ca^2+^-induced direct vesicle-to-hexosome colloidal transformation.

In the MAG-DHA/DOPG/α-tocopherol 56:30:14 (w/w/w) nanodispersion, which contained a lower DOPG fraction, corresponding to a DOPG/Ca^2+^ molar ratio of 3.9, an internal H_2_ phase was already detected when the aqueous buffer contained 3 mM Ca^2+^. This indicates that the direct lamellar (L_α_)-to-H_2_ phase transition occurred at a lower Ca^2+^ concentration than in the DOPG-richer MAG-DHA/DOPG/α-tocopherol 48:40:12 (w/w/w) nanodispersion. Clear H_2_ characteristic Bragg peaks were detected at *q* values of approximately 1.22, 2.10, and 2.42 nm^−1^, corresponding to a lattice parameter of approximately 5.98 nm. Further increasing the Ca^2+^ concentration to 5 mM in the aqueous buffer decreased the DOPG/Ca^2+^ ratio to 2.4. This was accompanied by a slight decrease in the lattice parameter of the internal H_2_ phase to approximately 5.73 nm, with corresponding Bragg peaks detected at *q* values of approximately 1.27, 2.19, and 2.53 nm^−1^. The different structural responses of the two nanodispersions upon exposure to Ca^2+^-containing buffer show that the Ca^2+^ concentration required to induce the direct vesicle-to-hexosome colloidal transformation depends strongly on lipid composition. The slight decrease in the H_2_ lattice parameter at higher Ca^2+^ concentration is consistent with the Ca^2+^-mediated dehydration and tighter packing of DOPG molecules embedded at the interfacial film, as discussed above. Our results are consistent with previous SAXS investigations of Ca^2+^-triggered lamellar-to-nonlamellar phase transitions in monoolein/DOPG nanodispersions, which demonstrated that Ca^2+^ promotes composition- and concentration-dependent structural transitions from weakly correlated bilayers or sponge (L_3_) phases toward inverse nonlamellar liquid crystalline self-assemblies, including the H_2_ phase^74,75^.

It is worth noting that, in the few available reports on the continuous microfluidic production of cubosomes and hexosomes, SAXS characterization has typically been performed on collected nanodispersions after microfluidic production rather than directly during their formation under flow conditions^79–85^. In some of these studies, additional post-processing steps were required before structural characterization. For example, Kim et al.^80^ reported that microfluidic mixing initially produced ethanol-in-water nanosized droplets, followed by rotary evaporation at 75 mbar and 57 °C, during which transformation to cubosomes occurred after approximately 14 min. In two other studies, collected nanodispersions were subjected to overnight vacuum drying for ethanol removal or concentrated by multiple centrifugal filtration steps to obtain cubosomes before SAXS analysis^79,83^. In this context, the present SAXS-on-chip study provides direct insight into the structural evolution of hexosomes during continuous microfluidic production under flow conditions, without requiring solvent removal or concentration of the collected nanodispersions prior to SAXS characterization.

### 3.3 CG-MD simulations of the Ca^2+^-induced Vesicle-to-Hexosome Transformation

To gain molecular-level insight into the experimentally observed Ca^2+^-induced direct L_α_-to-H_2_ phase transition, CG-MD simulations were performed using a multilamellar lipid model system composed of MAG-DHA/DOPG/α-tocopherol 48:40:12 (w/w/w) at a DOPG/Ca^2+^ molar ratio of 2. The simulations were carried out in triplicate using the Martini 3 CG force field^51^, which has previously been shown to capture L_α_-to-H_2_ phase transitions^54,86^ in several lipid systems.

After initial assembly, the triplicate simulations showed that, within approximately 2 µs, the multilamellar system evolved from an L_α_-like arrangement toward an H_2_-like organization. This transition proceeded through the formation of stalk-like intermediates, where adjacent bilayers came into close contact and subsequently reorganized into a continuous nonlamellar structure (**Figure 7A**). Visual inspection of the trajectories indicated that the first interbilayer contacts were primarily initiated by MAG-DHA molecules from opposing leaflets. These contacts expanded and reorganized into the final H_2_-like structure (**Figure 7A**, bottom snapshot), supporting the role of MAG-DHA in promoting negative interfacial curvature and nonlamellar phase formation.

**Figure 7.**
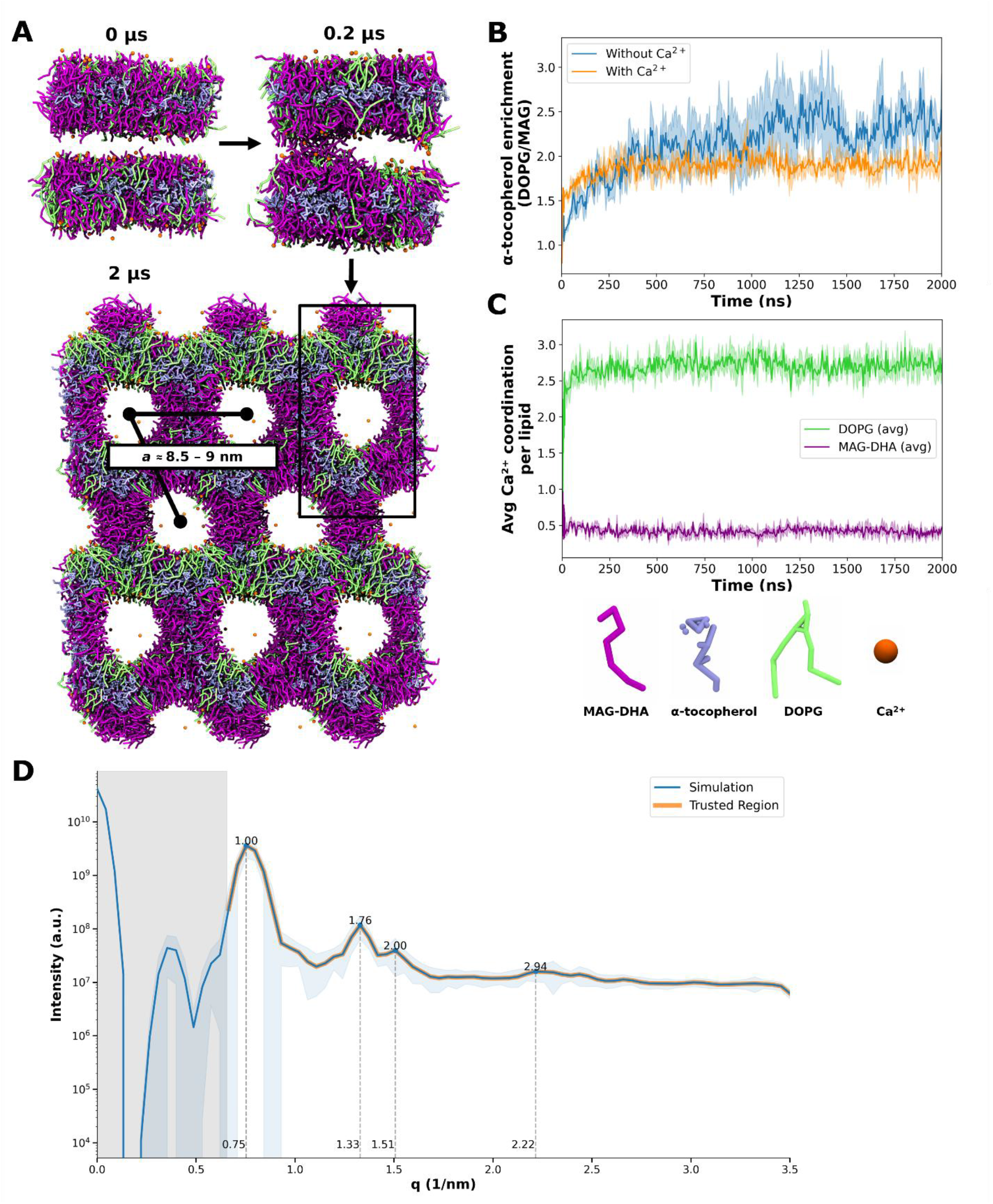
CG-MD simulations reveal how calcium ions drive the structural transformation from the lamellar phase to the inverted hexagonal (H_2_) phase. (A) Representative snapshots from CG-MD simulations illustrating the early stage of the lamellar-to-H_2_ phase transition in a multilamellar system composed of MAG-DHA/DOPG/α-tocopherol 48:40:12 (w/w/w) in the presence of Ca^2+^ ions. The process involves the formation of intermediate stalk-like structures as the bilayers come into contact (middle), and finally to the emergence of a H_2_ phase (bottom). In this latter stage, a clear spatial segregation between DOPG lipids (green) and MAG-DHA lipids (purple) is observed. (B) Time evolution of α-tocopherol enrichment at the DOPG/MAG-DHA interface in the absence (blue) and presence (orange) of Ca^2+^ ions. The preference was quantified by calculating the ratio of α-tocopherol –DOPG and α-tocopherol –MAG-DHA contacts within a 0.47 nm cutoff. (C) Temporal evolution of Ca^2+^ ions mean coordination with MAG-DHA and DOPG. Y axis values are average interaction of Ca^2+^ ions with each lipid species at a cutoff of 0.47 nm, over time. (D) SAXS profile obtained from CG-MD configurations for one representative replicate. The gray-shaded region indicates the low-*q* range excluded due to finite simulation-box and periodic-boundary effects. Within the reliable (*q*) range, reflections at approximately q values of 0.75, 1.33, and 1.51 nm^−1^ are consistent with the expected (10), (11), and (20) reflections of an H_2_ phase, respectively.

To clarify the role of Ca^2+^ ions during the L_α_-to-H_2_ phase transition, Ca^2+^–lipid interactions were quantified according to the procedure described in Section 2.8.2. Ca^2+^ ions preferentially coordinated with DOPG headgroups rather than with MAG-DHA (**Figure 7C**), consistent with the anionic nature of DOPG and its strong affinity for divalent cations^74,87^. On average, approximately 1.2 Ca^2+^ ions were coordinated per DOPG molecule. This preferential coordination supports the experimental interpretation that Ca^2+^–DOPG interactions promote dehydration of DOPG headgroups and tighter packing of DOPG molecules at the lipid–water interface, thereby facilitating the vesicle-to-hexosome transformation. Analysis of α-tocopherol distribution further showed preferential colocalization with DOPG (**Figure 7 Figure 7B**), particularly in the absence of Ca^2+^ ions, indicating that Ca^2+^–DOPG coordination modulates the local organization of α-tocopherol within the membrane. Together, these results suggest a cooperative mechanism in which Ca^2+^-DOPG coordination, the intrinsic H_2_-forming propensity of MAG-DHA, and α-tocopherol redistribution collectively support the transition toward the H_2_ phase.

The simulated transition can be rationalized by the molecular characteristics of the lipid components. MAG-DHA (**Figure 4**) behaves as a wedge-shaped amphiphile, with its highly unsaturated DHA chain providing conformational flexibility that supports the negative interfacial curvature required for H_2_ phase formation^77^. As discussed above, DOPG favors the L_α_ phase in excess water, whereas Ca^2+^ coordination with DOPG headgroups can reduce electrostatic repulsion and promote tighter interfacial packing. This Ca^2+^-mediated reduction in the effective headgroup area of DOPG can be rationalized using the critical packing parameter^88^, *CPP* = *v_s_*/(*a*_0_*l_c_*), where *v_s_* is the effective hydrophobic chain volume, *a*_0_ is the effective headgroup area of DOPG, and *l_c_* is the hydrophobic chain length. A reduction in *a*_0_, associated primarily with Ca^2+^ binding to DOPG headgroups and potentially supported by α-tocopherol redistribution within the interfacial area, increases the *CPP* value and, together with the intrinsic H_2_-forming propensity of MAG-DHA, promotes emergence of the internal H_2_ phase, corresponding to hexosome formation.

Control simulations were also performed to examine the composition dependence of L_α_ phase formation in the absence of Ca^2+^ ions. Although the SAXS pattern predominantly indicated L_α_ phase formation upon removal of Ca^2+^ ions for the MAG-DHA/DOPG/α-tocopherol 48:40:12 (w/w/w) composition (**Figure 6A**), this transition was not observed within the simulated timescale when the same composition was used in the CG-MD model. Instead, an L_α_ phase was obtained only in DOPG-enriched systems, either for the MAG-DHA/DOPG/α-tocopherol 34.3:56.6:9.2 (w/w/w) composition without Ca^2+^ ions (**Figure S5A**) or for the MAG-DHA/DOPG 37:63 (w/w) composition without α-tocopherol (**Figure S5B**). This indicates that L_α_ phase formation in the CG model is sensitive to the DOPG content, with the simulated phase transition occurring at a different lipid composition than indicated by the SAXS results (**Figure 6A**). Such composition-dependent shifts are commonly observed in coarse-grained simulations of lipid systems, including Martini-based models, in which qualitative phase behavior is often reproduced more reliably than exact experimental phase boundaries. For example, Pedersen et al. ^54^ showed that the reparametrized Martini 3 lipid models captured several features of the experimental DPPC/DOPC/cholesterol ternary phase diagram, although the simulated coexistence regions did not fully coincide with those experimentally determined.

In the present simulations, additional factors may have contributed to the difference between SAXS-derived and CG-MD phase behavior. The CG-MD systems did not include stabilizer molecules and were performed at lower water content than the experimental nanodispersions. In addition, the ethanol-containing solvent environment present during microfluidic mixing was not included in the CG-MD model. Ethanol can partition into lipid interfacial regions and alter lipid-chain ordering, membrane fluidity, and permeability,^72^ and may therefore affect the kinetics of lipid rearrangement. This simplification, together with the absence of stabilizer molecules, may contribute to the observed shift between the SAXS-derived and simulated phase behavior. Such simplifications are commonly used in computational studies of lipid self-assemblies^57,89^, to reduce system complexity and computational cost while still while retaining sufficient detail to capture the main structural features. The slight composition-dependent shift observed here may also arise from finite system size and the use of periodic boundary conditions, both of which are inherent to MD simulations. Nevertheless, the simulations reproduced the experimentally observed Ca^2+^- and composition-dependent trends.

Finally, a SAXS profile was calculated from a representative CG configuration to further support the structural assignment of the simulated Ca^2+^-containing system (**Figure 7D**). Additional SAXS profiles obtained by ensemble averaging configurations from the two independent replicas are presented in **Figure S6.** The low-(q) region below approximately 0.7 nm^−1^ was excluded from interpretation because it falls outside the reliable range accessible given the finite dimensions of the simulation box. Within the reliable *q* range, the calculated profile exhibited reflections at approximately *q* = 0.75, 1.33, and 1.51 nm^−1^. These reflections were assigned to the (10), (11), and (20) reflections of an H2 phase, respectively, with relative positions of 1:1.77:2.01, close to the expected hexagonal ratio of 1:√3:2. The lattice parameters calculated from the three reflections gave an average value of 9.58 ± 0.12 nm. Although larger than the experimental H_2_ lattice parameter of approximately 6 nm derived from the SAXS data (**Table 2**), the simulated value is qualitatively consistent with the more expanded hexagonal packing and larger aqueous channels observed in the CG structures. Minor deviations from the ideal H_2_ peak positions and from the experimental lattice parameter are expected and may arise from coarse graining, finite-size effects, and residual structural fluctuations within the simulated assembly. Overall, the calculated SAXS profile is consistent with the formation of an H_2_ phase and agrees with the structural evolution observed directly in the MD trajectories.

**Table 2.** SAXS-derived structural parameters and hydrodynamic size characteristics of MAG-DHA/DOPG/α-tocopherol nanodispersions prepared by microfluidic mixing at lipid compositions of 48:40:12 and 56:30:14 (w/w/w).^a^.

| Lipid System (wt%) | Ca<sup>2+</sup> Conc. | FRR | Nanodispersion | Space group | $a(nm)$ | Size (nm) ± SD | PDI ± SD |
| --- | --- | --- | --- | --- | --- | --- | --- |
| 48:40:12 | Ca<sup>2+</sup>-Free | 10 | Vesicles | L<sub>α</sub> | — | 125.7±1.4 | 0.20±0.02 |
| | | 20 | Vesicles | L<sub>α</sub> | — | 92.9±0.6 | 0.20±0.01 |
| | | 30 | Vesicles | L<sub>α</sub> | — | 79.9±0.3 | 0.20±0.01 |
| | 5mM | 10 | Hexosomes | H<sub>2</sub> | 6.21 | 174.4±0.9 | 0.18±0.02 |
| | | 20 | Hexosomes | H<sub>2</sub> | 5.95 | 129±0.9 | 0.17±0.01 |
| | | 30 | Hexosomes | H<sub>2</sub> | 5.99 | 124.2±0.8 | 0.18±0.01 |
| 56:30:14 | Ca<sup>2+</sup>-Free | 10 | Vesicles | L<sub>α</sub> | — | 148.2±0.5 | 0.22±0.01 |
| | | 20 | Vesicles | L<sub>α</sub> | — | 100.3±1.8 | 0.21±0.01 |
| | | 30 | Vesicles | L<sub>α</sub> | — | 83.1±0.4 | 0.20±0.01 |
| | 5mM | 10 | Hexosomes | H<sub>2</sub> | 5.73 | 133.2±1.0 | 0.14±0.01 |
| | | 20 | Hexosomes | H<sub>2</sub> | 5.6 | 126.5±1.0 | 0.16±0.01 |
| | | 30 | Hexosomes | H<sub>2</sub> | 5.64 | 130.9±0.7 | 0.16±0.01 |

<sup>a</sup>The lattice parameters of identified H<sub>2</sub> phases as derived from SAXS patterns presented in (Figure S8 A,B).

### 3.4. FRR- and Ca^2+^-Dependent Modulation of Nanodispersion Size and Internal Structure

To evaluate how microfluidic mixing conditions affected the size characteristics of the produced nanodispersions, DLS measurements were performed at FRRs of 10, 20, and 30 while maintaining a constant TFR of 200 μL min^−1^. All samples exhibited unimodal nanoparticle size distributions (**Figure S7**), and the corresponding hydrodynamic diameters (sizes) and polydispersity indices (PDIs) are summarized in **Table 2**.

In the absence of Ca^2+^ ions, both MAG-DHA/DOPG/α-tocopherol compositions, 48:40:12 and 56:30:14 (w/w/w), showed a pronounced FRR-dependent decrease in nanoparticle size. Increasing the FRR from 10 to 30 reduced the mean nanoparticle size from 125.7 ± 1.4 to 79.9 ± 0.3 nm for the 48:40:12 (MAG-DHA/DOPG/α-tocopherol, w/w/w) composition and from 148.2 ± 0.5 to 83.1 ± 0.4 nm for the 56:30:14 (MAG-DHA/DOPG/α-tocopherol, w/w/w) composition. Under these conditions, the vesicular nanodispersions showed moderate polydispersity, with PDI values around 0.20. This FRR-dependent nanoparticle size reduction is consistent with previous reports on microfluidic lipid nanoparticle production, where increasing the FRR enhances solvent exchange by narrowing the central ethanol-rich stream and strengthening solvent-concentration gradients between the ethanol-rich stream and the surrounding aqueous buffer ^84,90–92^. At higher FRRs, the accelerated solvent exchange likely promotes lipid self-assembly and colloidal closure into nascent vesicles, thereby limiting further nanoparticle growth and resulting in smaller nanoparticle sizes^15,81,91^.

In the presence of 5 mM Ca^2+^ ions in the aqueous buffer before microfluidic mixing, SAXS confirmed formation of hexosomes at all three investigated FRRs (**Table 2**). The lipid and ethanol concentrations of the produced nanodispersions are summarized in **Table S2**. For the MAG-DHA/DOPG/α-tocopherol 48:40:12 (w/w/w) composition, increasing the FRR from 10 to 30 decreased the DOPG/ Ca^2+^ molar ratio from 3.0 to 1.0 (**Table 1**) and was accompanied by a decrease in mean nanoparticles size from 174.4 ± 0.9 to 124.2 ± 0.8 nm (**Table 2**). The lattice parameter of the internal H_2_ phase decreased from 6.21 nm at FRR 10 to 5.95 nm at FRR 20 and then remained nearly unchanged at FRR 30 (5.99 nm) (**Table 2**). This lattice-parameter evolution is consistent with increased Ca^2+^ local concentration at higher FRR, favoring Ca^2+^-DOPG association, dehydration of DOPG headgroups, and tighter packing of DOPG molecules within at the lipid–water interfacial film, as discussed above. The relatively larger lattice parameter observed at FRR 10 may also be influenced by the higher residual ethanol content (4.18 wt%) in the nanodispersion. These SAXS results are in line with previous studies in which ethanol-induced increases in lattice parameters were attributed to partial partitioning of ethanol into the hydrophilic domains of internal hydrophilic domains of nonlamellar liquid crystalline nanostructures^12,82,93^. For instance, an ethanol concentration-dependent increase in the lattice parameter of the internal cubic Pn3m phase was reported for Pluronic F127-stabilized phytantriol cubosomes^94^. In contrast, the lower ethanol contents at FRR 20 and 30 (2.18 and 1.47 wt%, respectively), together with the lower DOPG/Ca^2+^ molar ratios, are consistent with the formation of more compact internal H_2_ structures. However, further reduction in the lattice parameter of the internal H_2_ phase appears limited at the lowest DOPG/Ca^2+^ ratio.

For the MAG-DHA/DOPG/α-tocopherol 56:30:14 (w/w/w) composition, the mean sizes of the produced hexosomes showed only minor variation across the investigated FRR range and remained within 126-133 nm, with reduced PDI values reaching 0.14, indicating relatively narrow nanoparticle size distributions (**Table 2**). The corresponding lattice parameters of the internal H_2_ phases were consistently smaller than those observed for the MAG-DHA/DOPG/α-tocopherol 48:40:12 composition (**Table 2**). This is consistent with the lower DOPG fraction in this composition, which likely reduces electrostatic repulsion among DOPG molecules embedded at the lipid-water interface. In line with previous reports^76,95,96^ on charged amphiphile-containing lyotropic liquid crystalline (LLC) nanodispersions, this reduced repulsion is associated with lower water uptake within the water nanochannels of the internal H_2_ phase, resulting in smaller lattice parameters.

### 3.5. Morphological Characterization of Liquid Crystalline Nano-Self-Assemblies

In addition to SAXS analysis and DLS-based nanoparticle size characterization, cryo-TEM was employed as a complementary technique to visualize the morphological features of three selected nanodispersions prepared in the absence and presence of Ca^2+^ ions. Such complementary imaging is particularly relevant for nonlamellar liquid crystalline nanodispersions, which often exhibit intrinsic structural heterogeneity^76,78^. In particular, the incorporation of phospholipids may increase the fraction of coexisting vesicular nanostructures.

For the MAG DHA/DOPG/α-tocopherol (56:30:14, w/w/w) nanodispersion prepared at FRR 10 and TFR = 200 µL min^−1^ using Ca^2+^-free PIPES buffer (10 mM, pH 7.0), representative cryo-TEM micrographs (**Figure 8 A,B**) revealed a dominant population of relatively small, spherical vesicular nanostructures, mainly unilamellar and oligolamellar vesicles (ULVs and OLVs) with diameters in the range of approximately 8–60 nm (marked with blue arrows). A small fraction of larger spherical vesicular nanoobjects, including ULVs and bilamellar vesicles (BLVs), was also detected, with diameters of around 90–160 nm (marked with yellow arrows in **Figure 8 A,B**). These observations are consistent with the corresponding SAXS patterns (**Figure 6 B**), which showed diffuse scattering with broad low-*q* correlation peaks, characteristic of weakly correlated bilayers.

**Figure 8.**
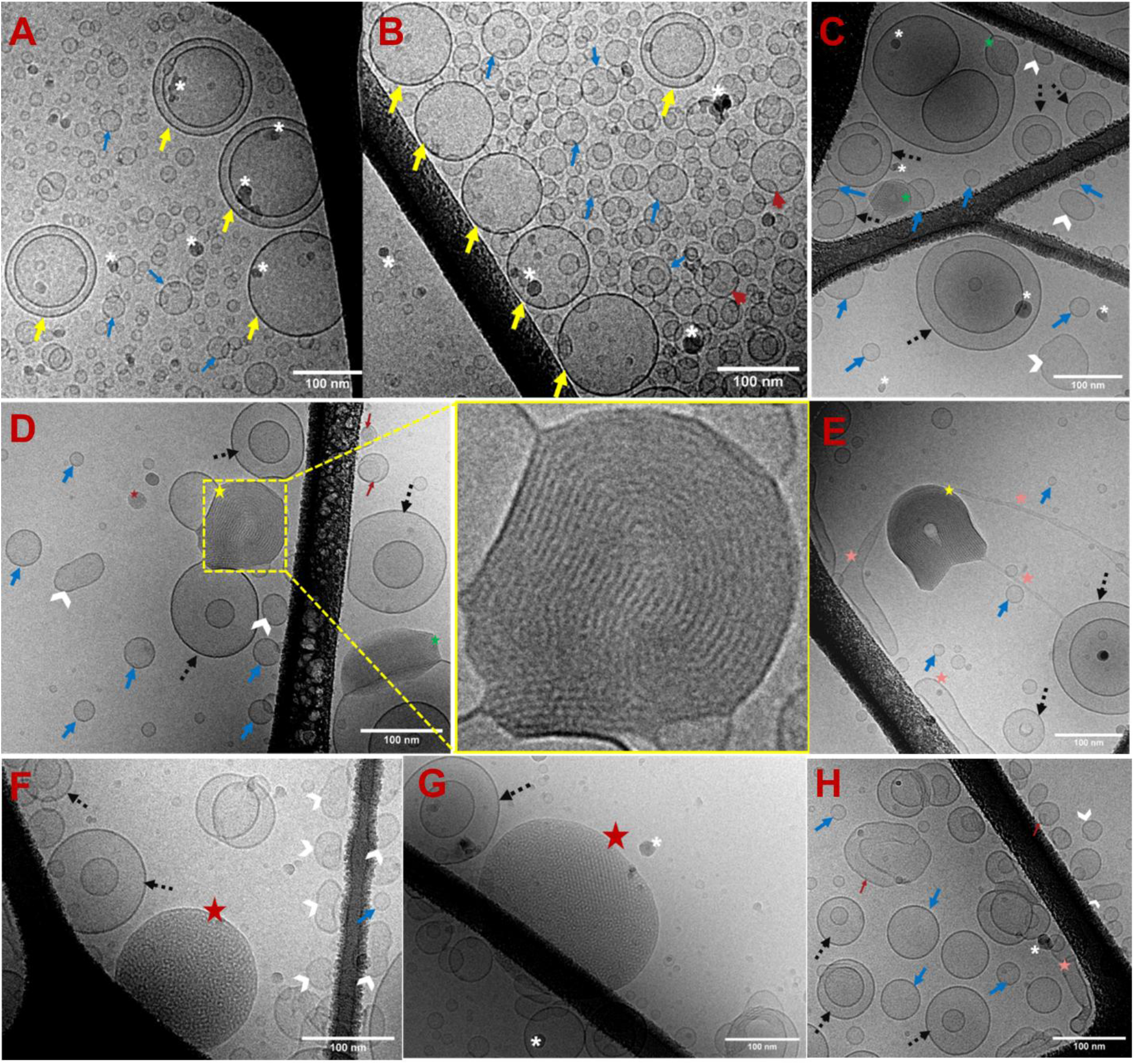
Representative cryo-TEM micrographs of selected MAG-DHA/DOPG/α-tocopherol nanodispersions prepared in the absence and presence of Ca^2+^ ions. Panels (A,B) show the nanodispersion prepared at a MAG-DHA/DOPG/α-tocopherol weight ratio of 56:30:14, FRR 10, and TFR = 200 µL min^−1^ using Ca^2+^-free PIPES buffer (10 mM, pH 7.0). Relatively small spherical unilamellar and oligolamellar vesicles (ULVs and OLVs) are marked with blue arrows, whereas larger vesicular nanoobjects, including ULVs and bilamellar vesicles (BLVs), are marked with yellow arrows. Panels (C–E) show the corresponding MAG-DHA/DOPG/α-tocopherol (56:30:14, w/w/w) nanodispersion prepared in the presence of 5 mM Ca^2+^ ions. Hexosomes displaying characteristic internal curved striations are marked with yellow stars and are more clearly visible in the inset of panel (D). Coexisting ULVs and OLVs are marked with blue arrows and dashed black arrows. Distorted vesicular nanoobjects, folded vesicles, multicompartment nanoobjects, and elongated tubular-like vesicular structures are marked with white arrowheads, red arrows, green stars, and pink stars, respectively. Panels (F–H) show the MAG-DHA/DOPG/α-tocopherol nanodispersion prepared at a weight ratio of 48:40:12, FRR = 20, and TFR = 200 µL min^−1^ in the presence of 3 mM Ca^2+^ ions. Relatively large nanoobjects with dense internal features, most likely hexosomes, are marked with red stars. Coexisting ULVs and OLVs are marked with blue arrows and dashed black arrows, whereas distorted and elongated vesicular nanoobjects are marked with white arrowheads. Scale bars correspond to 100 nm.

Upon the introduction of 5 mM Ca^2+^ ions in the aqueous buffer, pronounced morphological alterations were observed (**Figure 8 C–E**). Notably, nanoparticles displaying internal curved striations typical of hexosomes, were detected particularly in panels (D) and (E), and in the inset of panel (D) in Figure 8 (marked with yellow stars), supporting the SAXS-based assignment of the internal H_2_ phase^49^. In addition to hexosomes, ULVs and OLVs with diameters in the ranges of approximately 10–35 nm and 70–178 nm were still observed (marked with blue and dashed black arrows in **Figure 8 C–E**, respectively). Moreover, nanoobjects with distorted structure (mark with white arrowheads), folded vesicles (mark with red arrows) and multicompartment nanoobjects (marked with green stars) were detected. Elongated vesicular nanoobjects were also observed, including long tubular vesicles and thin line-like vesicular nanostructures connecting adjacent nanoparticles (marked with pink star in **Figure 8 E**). These morphologies are reminiscent of block liposomes, for which connected nanoscale spheres, pears, tubes, and rods have been reported in membranes containing charged lipid components^97^. However, a contribution from the microfluidic mixing conditions to the formation of such tubular or rod-like vesicular nanostructures cannot be excluded. The coexistence of these morphologically distinct nanoobjects suggests that the Ca^2+^-triggered structural transition is associated with pronounced morphological heterogeneity and does not proceed uniformly across the entire nanoparticle population.

Cryo-TEM investigations were also conducted on the MAG-DHA/DOPG/α-tocopherol (48:40:12, w/w/w) nanodispersion prepared at FRR 20 and TFR 200 µL min^−1^ in the presence of 3 mM Ca^2+^ ions in the aqueous buffer (**Figure 8F–H**). The cryo-TEM micrographs showed nanoparticles with dense internal features, marked with red stars in **Figure 8 F,G**. These nanoobjects are most likely hexosomes based on the SAXS analysis (**Figure 6A**), although their dense internal features did not allow unambiguous structural assignment by cryo-TEM alone. In addition, coexisting ULVs and OLVs with sizes in the range of 15–110 nm were detected (marked with blue and black dashed arrows in **Figure 8 F–H**, respectively). Furthermore, a relatively small distorted and long vesicular structure, including tubular-like vesicular nanoobjects, was also detected, as marked with white arrowheads. The presence of such vesicular nanoobjects is in line with previous cryo-TEM studies of DOPG- and other charged lipid-containing nonlamellar liquid crystalline nanodispersions, where relatively small coexisting vesicular structures were observed^49,76,78^.

## 4. CONCLUSIONS

This work demonstrates, to the best of our knowledge for the first time, the use of a 3D-printed X-ray-compatible microfluidic chip for online synchrotron SAXS-on-chip monitoring of Ca^2+^-triggered hexosome formation during continuous microfluidic production. In contrast to previous studies relying mainly on off-chip SAXS analysis of produced nonlamellar liquid crystalline nanodispersions typically cubosomes and hexosomes^79–85^, where SAXS investigations were mainly performed on nanodispersions following their preparation^79–85^, the present platform enabled direct structural characterization inside the microfluidic channel and under flow conditions, thereby avoiding ethanol removal or other post-processing steps prior to SAXS measurements.

By integrating online synchrotron SAXS-on-chip with *ex situ* SAXS measurements of collected nonlamellar liquid crystalline nanodispersions, together with DLS, cryo-TEM, and CG-MD simulations, this study links microfluidic process conditions to the structural, size, and morphological evolution of MAG-DHA/DOPG/α-tocopherol nano-self-assemblies. The online SAXS-on-chip findings indicated rapid Ca^2+^-induced formation of hexosomes on a tens-of-milliseconds time scale during mixing of the ethanolic lipid solution with the Ca^2+^-containing aqueous medium, with the first three Bragg reflections characteristic of the H_2_ phase were detected at the earliest probed position. For the MAG-DHA/DOPG/α-tocopherol 48:40:12 (w/w/w) composition, the initially detected biphasic H_2_ feature evolved toward a neat internal H_2_ phase after approximately 14 ms of residence time, whereas the corresponding MAG-DHA/DOPG/α-tocopherol 56:30:14 (w/w/w) composition retained the biphasic H_2_ feature at the investigated channel positions within the microfluidic channel, highlighting that the internal H_2_ structure is sensitive to lipid composition.

At the two investigated MAG-DHA/DOPG/α-tocopherol compositions, the *ex situ* SAXS findings identify the DOPG/Ca^2+^ molar ratio as a key parameter directing the Ca^2+^-triggered vesicle-to-hexosome transformation. Lowering this ratio induced colloidal transformation from vesicular nanodispersions to hexosomes and yielded more compact H₂ nanostructures, consistent with Ca^2+^-mediated dehydration and tighter packing of DOPG headgroups at the lipid–water interface. This interpretation is supported by CG-MD simulations, which showed preferential Ca^2+^ coordination with DOPG, MAG-DHA-driven negative interfacial spontaneous curvature, and Ca^2+^-dependent redistribution of α-tocopherol within the interfacial area. Together with DLS and cryo-TEM, these findings show that Ca^2+^-triggered hexosome formation is accompanied by changes in nanoparticle size and pronounced morphological heterogeneity, including coexistence of hexosomes and vesicular nanostructures.

Collectively, these findings demonstrate the utility of the 3D-printed X-ray-compatible microfluidic chip as a proof-of-concept platform for position-resolved online SAXS-on-chip investigations of Ca^2+^-triggered hexosome formation during continuous microfluidic production. By collecting SAXS patterns at defined channel positions, the platform enabled direct probing of the evolving internal H_2_ structure under flow, prior to off-chip collection and ethanol removal. Future extensions of this approach will focus on drug-loaded cubosomes and hexosomes, where lipid composition, microfluidic channel geometry, residence time, mixing efficiency, and Ca^2+^ ion exposure can be directly related to the resulting internal nanostructure and nanoparticle morphology during production. Further development of microfluidic chip designs, including refined liquid handling, controlled microscale mixing, and, where appropriate, size-exclusion or fractionation elements, may help reduce the coexistence of vesicular and micellar structures with cubosomes and hexosomes, thereby helping to address the intrinsic heterogeneity of nonlamellar liquid crystalline nanodispersion^78^.

## Author Contributions

Z.B. led the experimental work, including methodology development, fabrication, data curation, formal analysis, and visualization, and prepared the original manuscript draft. M.V. performed the molecular dynamics simulations, including system setup and analysis, and contributed to manuscript revision. F.S. carried out MD data analysis and contributed to manuscript revision. M.D. contributed to supervision, experimental investigation, and formal data analysis. B.R. contributed to supervision and investigation. W.P. and P.C.T.S. conceptualized the molecular dynamics simulations, secured funding, supervised the work, and contributed to manuscript revision. S.S.K. contributed to supervision, provided research resources, and participated in manuscript revision. W.E.S. contributed to supervision and manuscript revision. A.Y. conceived the study, secured funding, supervised the project, curated data, and contributed to both the original draft and manuscript revision.

## ACKNOWLEDGMENT

Financial support was provided by the Novo Nordisk Foundation (Grant No. NNF22OC0079752 to A.Y.). A.Y. further acknowledges financial support from the Danish Council for Independent Research | Technology and Production Sciences (Reference DFF-3105-00039B) and the Danish Natural Sciences Research Council (DanScatt) for SAXS experiments. P.C.T.S. acknowledges support from the French National Centre for Scientific Research (CNRS). M.V. and P.C.T.S. acknowledge the PSMN (Pôle Scientifique de Modélisation Numérique) and the Centre Blaise Pascal’s IT test platform at ENS de Lyon, Lyon, France, for providing computational resources. The platform operates the SIDUS solution developed by Emmanuel Quemener ^98^. P.C.T.S. further acknowledge support from Sanofi. W.P. acknowledges funding from the Novo Nordisk Foundation (Grant Nos. NNF18SA0035142 and NNF22OC0079182). The authors gratefully acknowledge the use of the University of Copenhagen Small-Angle X-ray Scattering facility (CPHSAXS), funded by the Novo Nordisk Foundation (Grant No. NNF19OC0055857). Synchrotron SAXS data were collected at beamline P12, operated by EMBL Hamburg at the PETRA III storage ring, DESY, Hamburg, Germany. The authors acknowledge PolyFabLab, a research infrastructure at DTU Nanolab financially supported by the Novo Nordisk Foundation (Grant No. NNF21OC0068814), where microfluidic chip fabrication was carried out. The authors thank Dr. Clement Blanchet and Dr. Dmytro Soloviov for valuable support and technical assistance during the beamline experiments.

## ABBREVIATIONS

2D: two-dimensional
3D: three-dimensional
A: lattice parameter
BMF: Boston Micro Fabrication
CG: coarse-grained
CG-MD: coarse-grained molecular dynamics
CPP: critical packing parameter
cryo-TEM: cryogenic transmission electron microscopy
DHA-MAG: docosahexaenoic acid monoacylglycerol
DLS: dynamic light scattering
DOPG: dioleoylphosphatidylglycerol
FRR: flow-rate ratio
FWHM: full width at half maximum
HFF: hydrodynamic flow focusing
HTL: high-temperature resin
LLC: lyotropic liquid crystalline
MD: molecular dynamics
PDB: Protein Data Bank
PDI: polydispersity index
PµSL: projection microstereolithography
SAXS: small-angle X-ray scattering
SANS: small-angle neutron scattering
SEM: scanning electron microscopy
TFR: total flow rate.

