## Supplementary Information for "3D Printed X-ray Compatible Microfluidics for Online Characterization of Hexosomes: A Synchrotron SAXS-on-Chip Study with Molecular Dynamics Insights"

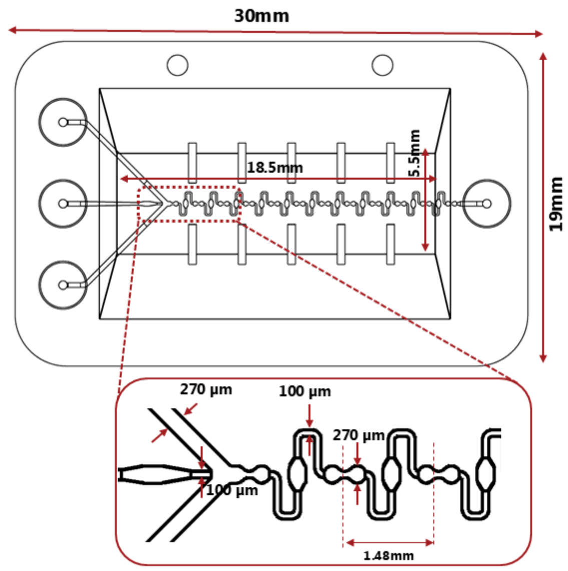


**Figure S 1.** Schematic illustration of the designed microfluidic system with alternating width of 270 µm and 100 µm system for online SAXS-on-chip investigations.

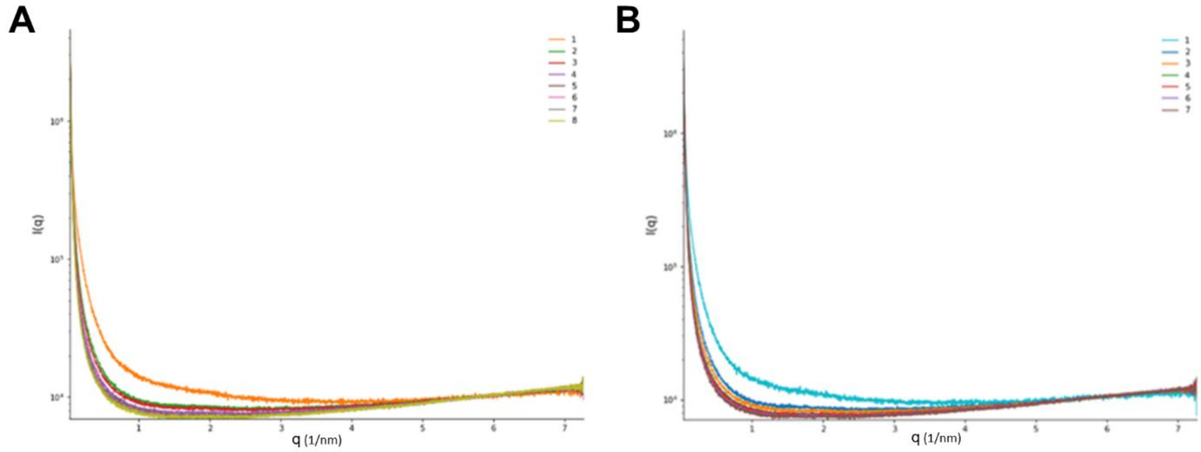


**Figure S 2.** Representative background SAXS patterns measured with PIPES buffer (10 mM, pH 7.0) in the central microfluidic channel at two selected positions: (A) position 1, before mixing, and (B) position 2, at the mixing point. The microfluidic chip was used without additional post-print ultraviolet curing. Minor changes in the background scattering were observed during the first 7–8 X-ray exposures, consistent with beam-induced changes in the resin, potentially involving further polymerization or crosslinking. After repeated exposures, the background scattering stabilized and became reproducible.

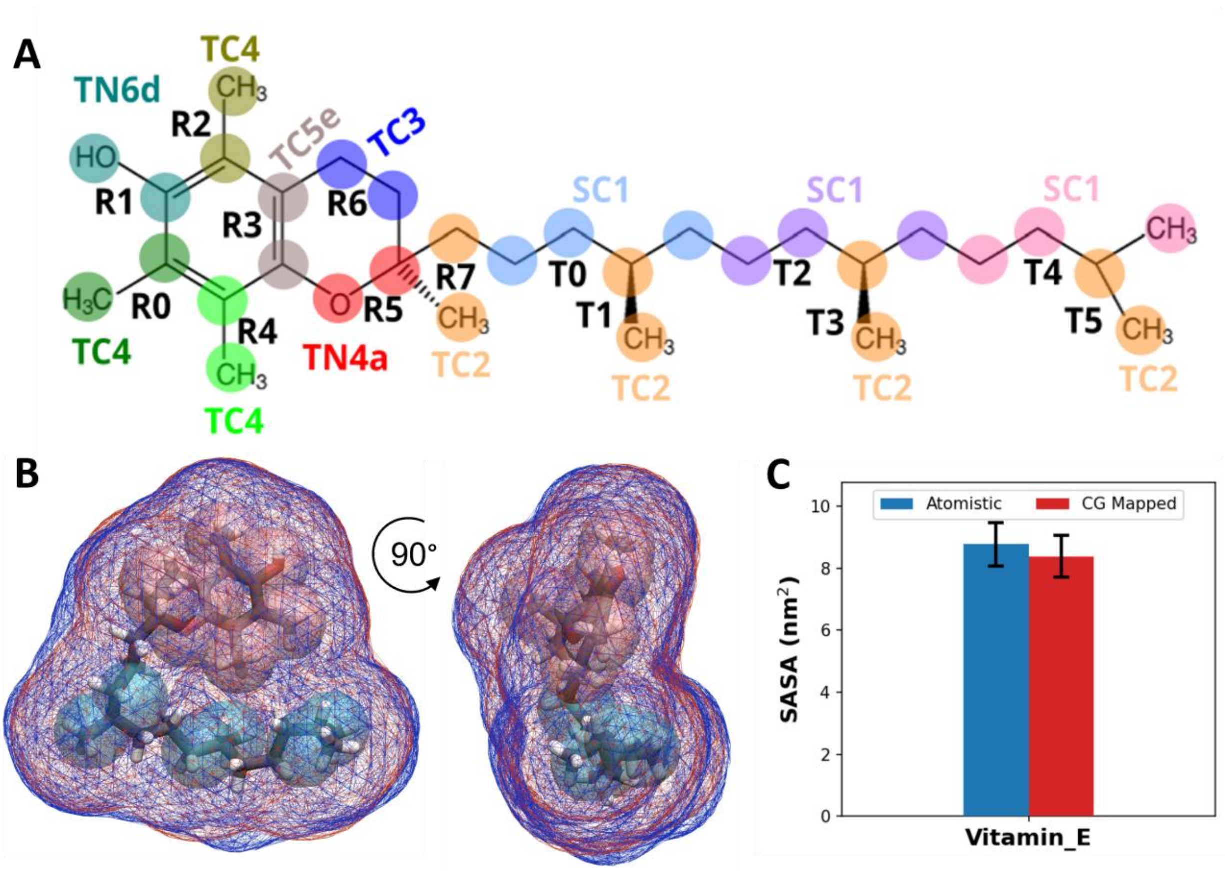


**Figure S 3.** α-tocopherol CG model description (A) Coarse-grained mapping over the atomistic structure. The colored circles identify the atoms to which the respective CG beads were mapped; the bead type is also indicated in a label of the same color. (B) Connolly surfaces for α-tocopherol from CG (red wireframes) and all-atom simulations (blue wireframes) (C) Mean solvent accessible surface area (SASA), with error bars corresponding to standard deviations, for CG-mapped (in red) and all-atom simulations (in blue).

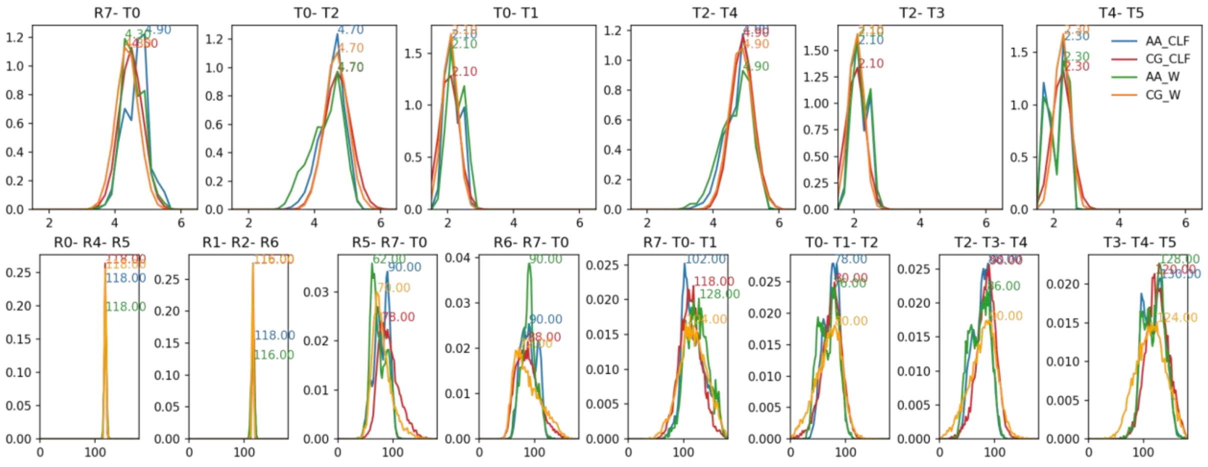

**Figure S 4.** Bonded distributions for α-tocopherol in atomistic and CG detail both in water and in chloroform.

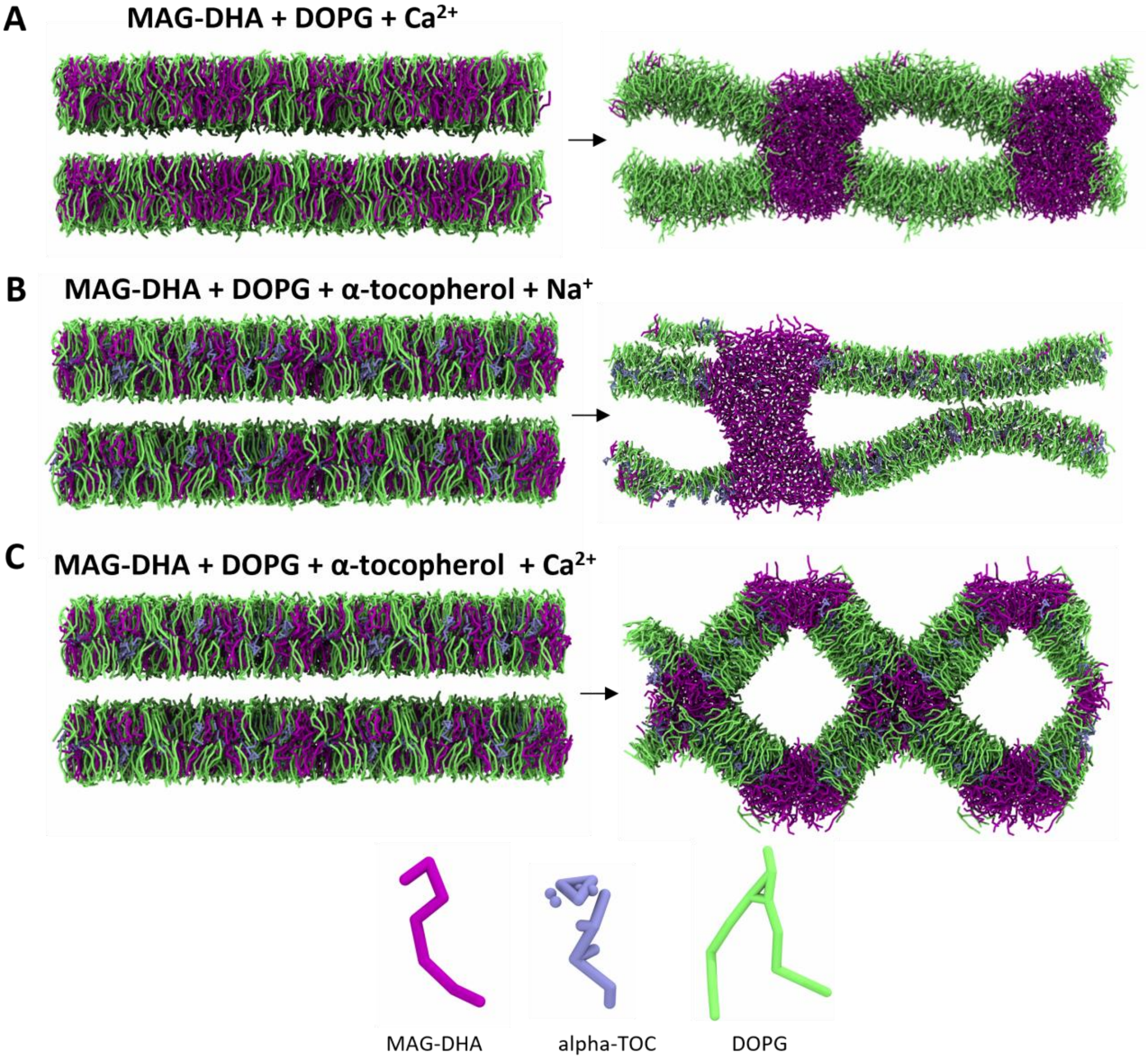


**Figure S 5.** MD snapshots representative frames of the beginning and end of the multilamellar system simulations. (A) MAG-DHA:DOPG (37:63, w/w) in the presence of $Ca^{2+}$ ions. (B) MAG-DHA:DOPG:α-tocopherol (34:57:9, w/w/w) in the absence of $Ca^{2+}$ ions. (C) MAG-DHA:DOPG:α-tocopherol (34:57:9, w/w/w) in the presence of $Ca^{2+}$ ions.

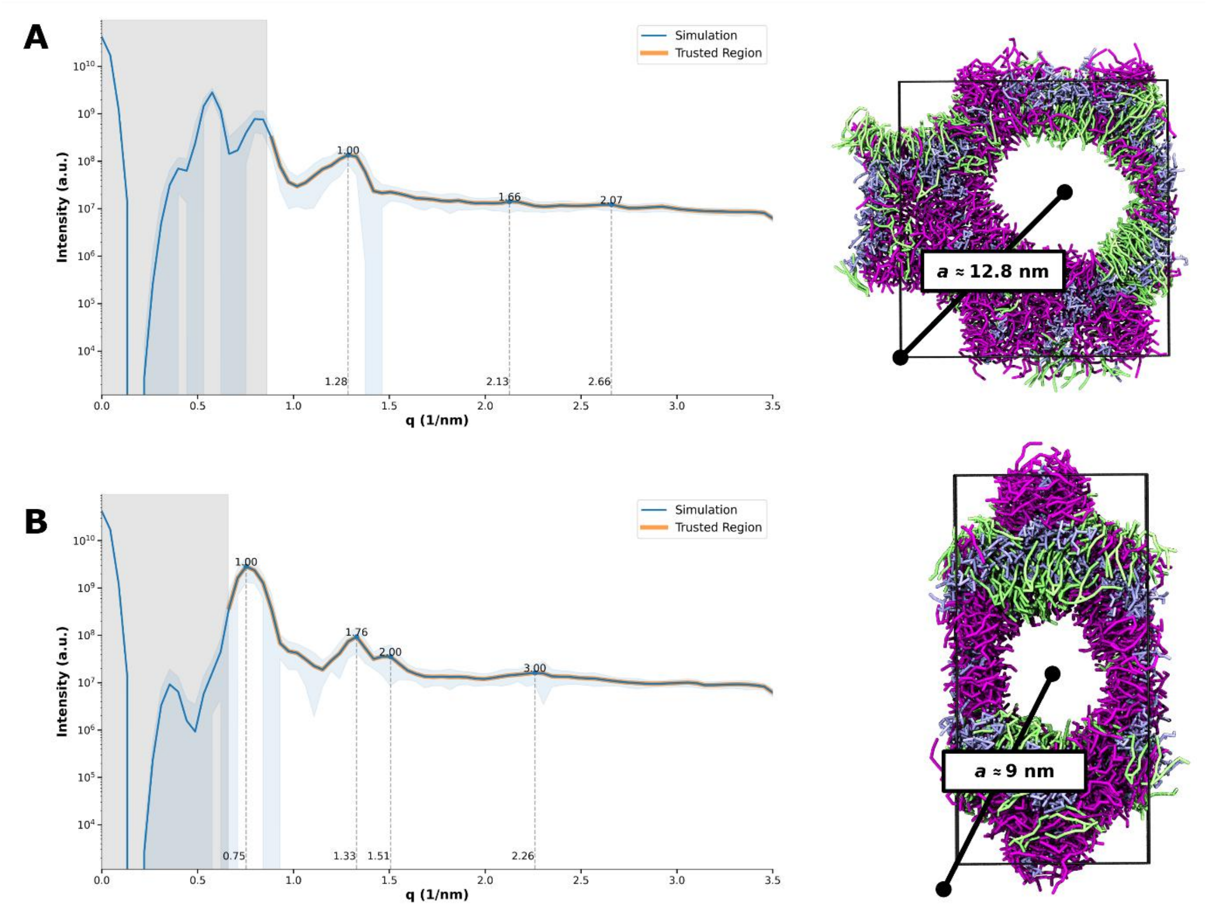


**Figure S6.** Simulated SAXS profiles generated from ensemble average of 100 frames over the last 500 ns of one of two of the replicas with a step size of 50 (A and B, left side). Blue curves show the simulated scattering intensity, while orange curves indicate the peak positions identified within the trusted q-range. Representative MD snapshots (right) show a single $H_2$ unit cell, with the center-to-center water channel spacing indicated for each structure. Replica B exhibits Bragg reflections consistent with the expected $H_2$ symmetry. In contrast, although replica A clearly displays an inverse hexagonal morphology, its larger water channels result in a larger lattice parameter, shifting the first Bragg reflection into the untrusted $q$ region affected by finite simulation box size. Consequently, the peak assignment algorithm cannot reliably identify the first-order reflection.

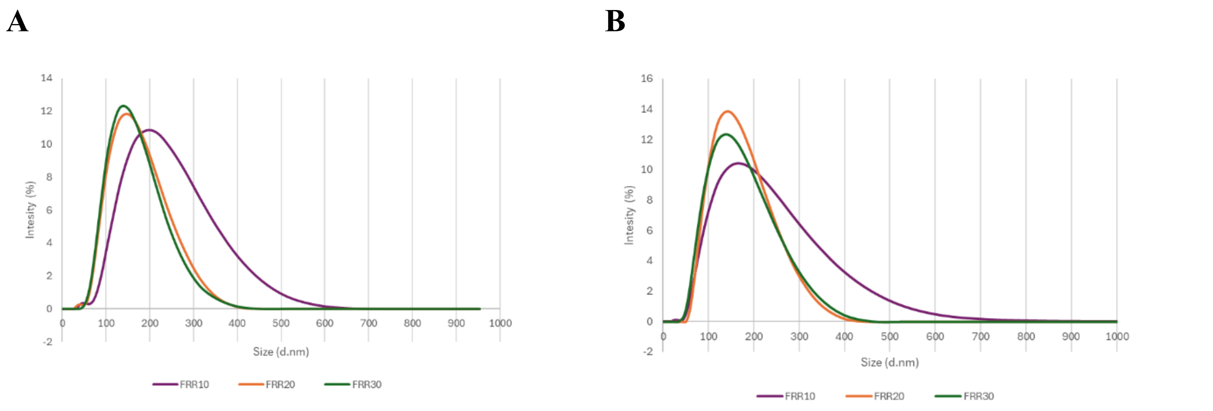


**Figure S 7.** DLS-derived particle size distributions of MAG-DHA/DOPG/α-tocopherol nanodispersions produced by microfluidic mixing in 5 mM $Ca^{2+}$-containing PIPES buffer at lipid compositions of 48:40:12 (A) and 56:30:14 (B) (w/w/w), using different FRRs of 10, 20, and 30 and a constant TFR of 200 μL $min^{-1}$.

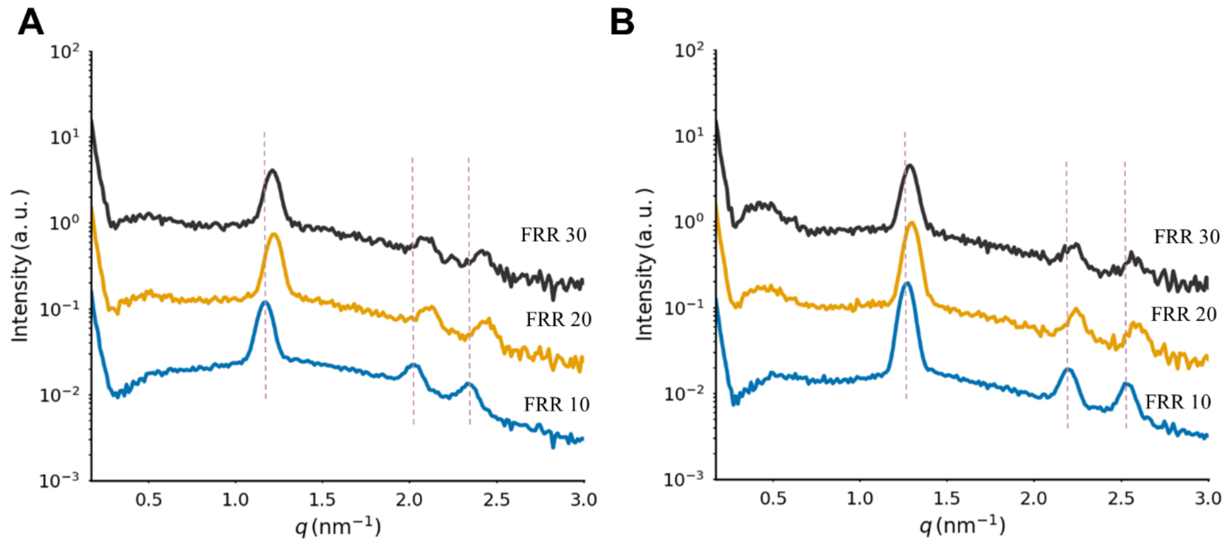


**Figure S 8.** SAXS profiles of MAG-DHA/DOPG/α-tocopherol nanodispersions prepared by microfluidic mixing in 5 mM $Ca^{2+}$-containing PIPES buffer at lipid compositions of 48:40:12 (A) and 56:30:14 (B) (w/w/w). The flow rate ratio (FRR) varied from 10 to 30 at a constant total flow rate (TFR) of 200 μL $min^{-1}$.

**Table S 1.** SAXS-derived structural parameters of MAG-DHA/DOPG/α-tocopherol nanodispersions along the microfluidic chip. The first hexagonal ($H_2$) phase is marked with blue dashed arrows in **Figure 4 A,B**, whereas the second coexistence $H_2$ phase is marked with red asterisks. $t_{res}$denotes the residence time downstream of the first mixing point; a is the lattice parameter of the $H_2$ phase.

| (MAG-DHA/DOPG/ α-tocopherol) | Position on chip | $t_{res}$ (ms) | $q$ Peaks ($nm^{-1}$) | Assigned Phase(s) | Lattice Parameter $a$ (nm) |
|---|---|---|---|---|---|
| 48:40:12 | 1 | Before mixning | - | Micellar sloution | - |
| | 2 | ~0 | 1.22, 2.11, 2.44 | $H_2$ (First) | 5.95 |
| | 2 | ~0 | 1.11, 1.92 | $H_2$ (Second) | 6.54 |
| | 3 | ~7 | 1.24, 2.15, 2.48 | $H_2$ (First) | 5.85 |
| | 3 | ~7 | 1.10, 1.92 | $H_2$ (Second) | 6.57 |
| | 4 | ~14 | 1.17, 2.04, 2.37 | $H_2$ | 6.15 |
| 56:30:14 | 1 | Before mixing | - | Micellar sloution | - |
| | 2 | ~0 | 1.27, 2.19 | $H_2$ (First) | 5.72 |
| | 2 | ~0 | 1.07, 1.86 | $H_2$ (second) | 6.77 |
| | 3 | ~7 | 1.29, 2.24, 2.59 | $H_2$ (First) | 5.61 |
| | 3 | ~7 | 1.19, 2.06 | $H_2$ (Second) | 6.1 |
| | 4 | ~14 | 1.28, 2.23, 2.58 | $H_2$ (First) | 5.64 |
| | 4 | ~14 | 1.21, 2.10 | $H_2$ (Second) | 5.98 |

**Table S 2.** Final lipid composition weight percentages calculated using python script below at different flow rate ratios (FRR).

| Lipid System (wt%) | FRR (%) | MAG-DHA (%) | DOPG (%) | α-tocopherol (%) | EtOH (%) | Water (%) | PIPES (%) | F127 (%) |
|---|---|---|---|---|---|---|---|---|
| | 10 | 1.34 | 1.12 | 0.33 | 4.18 | 1.12 | 91 | 0.92 |
| 48:40:12 | 20 | 0.7 | 0.58 | 0.17 | 2.18 | 0.58 | 94.83 | 0.96 |
| | 30 | 0.47 | 0.39 | 0.12 | 1.47 | 0.39 | 96.18 | 0.97 |
| | 10 | 1.61 | 0.86 | 0.40 | 4.31 | 0.86 | 91 | 0.92 |
| 56:30:14 | 20 | 0.84 | 0.45 | 0.21 | 2.27 | 0.45 | 94.85 | 0.96 |
| | 30 | 0.57 | 0.30 | 0.14 | 1.52 | 0.30 | 96.19 | 0.97 |

**Supplementary Code S1.** Python script used for calculating final compositions and lipids-to-$Ca^{2+}$ molar ratios.

```
"""
This script calculates the final composition (% w/w) of microfluidically
prepared MAG-DHA/DOPG/alpha-tocopherol nanodispersions obtained by mixing an
ethanol-lipid center stream with a calcium-containing aqueous buffer stream.

The script also calculates:
  - Final Ca2+ concentration after microfluidic mixing
  - DOPG/Ca2+ molar ratio
  - (DOPG + MAG-DHA)/Ca2+ molar ratio
"""

def calculate_final_percentages(FRR=10, TFR=200, calcium_conc_mM=5):
    """
    Calculate final component mass percentages and lipid-to-Ca2+ molar ratios.

    Parameters
    ----------
    FRR : float
        Flow rate ratio, defined as buffer stream : ethanol-lipid stream.

    TFR : float
        Total flow rate, in uL/min.

    calcium_conc_mM : float
        Ca2+ concentration in the buffer stream before mixing, in mM.

    Returns
    -------
    dict
        Dictionary containing the final Ca2+ concentration, component
        compositions (% w/w), and lipid-to-Ca2+ molar ratios.
    """
```

```python
# ---------------------------------------------------------------------
# Input formulation parameters
# ---------------------------------------------------------------------
lipid_fraction = 0.40       # Lipid fraction in ethanol-lipid stream, w/w
ethanol_fraction = 0.60     # Ethanol fraction in ethanol-lipid stream, w/w

lipid_composition = {
    "MAG-DHA": 0.48,        # Mass fraction of total lipid
    "DOPG": 0.40,           # Mass fraction of total lipid
    "alpha-tocopherol": 0.12,
}

# ---------------------------------------------------------------------
# Physical constants
# ---------------------------------------------------------------------
ethanol_density = 0.79      # mg/uL
buffer_density = 1.00       # mg/uL

dopg_mw = 797.00            # mg/mmol
mag_dha_mw = 402.57         # mg/mmol

# ---------------------------------------------------------------------
# Flow rates
# ---------------------------------------------------------------------
center_flow = TFR / (FRR + 1)       # Ethanol-lipid stream, uL/min
buffer_flow = TFR - center_flow     # Aqueous buffer stream, uL/min

# Ca2+ is assumed to be present only in the aqueous buffer stream.
final_calcium_mM = calcium_conc_mM * (buffer_flow / TFR)

# ---------------------------------------------------------------------
# Mass-balance scaling factor
# ---------------------------------------------------------------------
dopg_fraction = lipid_composition["DOPG"]
milliq_term = lipid_fraction * dopg_fraction

X = center_flow / (
    lipid_fraction + ethanol_fraction / ethanol_density + milliq_term
)

# ---------------------------------------------------------------------
# Mass contributions, mg/min
# ---------------------------------------------------------------------
lipid_total = lipid_fraction * X
ethanol = ethanol_fraction * X
milliq_water = dopg_fraction * lipid_total

buffer_mass = buffer_flow * buffer_density * 0.99
f127 = buffer_flow * buffer_density * 0.01

lipids = {
    component: fraction * lipid_total
    for component, fraction in lipid_composition.items()
}

total_mass = lipid_total + ethanol + milliq_water + buffer_mass + f127

# ---------------------------------------------------------------------
# Final formulation composition, % w/w
# ---------------------------------------------------------------------
results = {
    "Final Ca2+ concentration (mM)": final_calcium_mM,
    "MAG-DHA (% w/w)": lipids["MAG-DHA"] / total_mass * 100,
    "DOPG (% w/w)": lipids["DOPG"] / total_mass * 100,
    "alpha-tocopherol (% w/w)": lipids["alpha-tocopherol"] / total_mass * 100,
    "Ethanol (% w/w)": ethanol / total_mass * 100,
    "Milli-Q water (% w/w)": milliq_water / total_mass * 100,
```

```python
        "Buffer (% w/w)": buffer_mass / total_mass * 100,
        "F127 (% w/w)": f127 / total_mass * 100,
        "Total lipid (% w/w)": lipid_total / total_mass * 100,
    }

    # ------------------------------------------------------------------
    # Lipid-to-Ca2+ molar ratios
    # ------------------------------------------------------------------
    calcium_mmol = (buffer_flow * calcium_conc_mM) / 1_000_000

    dopg_mmol = lipids["DOPG"] / dopg_mw
    mag_dha_mmol = lipids["MAG-DHA"] / mag_dha_mw

    if calcium_mmol > 0:
        results["DOPG/Ca2+ (mol/mol)"] = dopg_mmol / calcium_mmol
        results["(DOPG + MAG-DHA)/Ca2+ (mol/mol)"] = (
            dopg_mmol + mag_dha_mmol
        ) / calcium_mmol
    else:
        results["DOPG/Ca2+ (mol/mol)"] = "N/A"
        results["(DOPG + MAG-DHA)/Ca2+ (mol/mol)"] = "N/A"

    return results


# ----------------------------------------------------------------------
# Example calculation
# ----------------------------------------------------------------------
if __name__ == "__main__":

    results = calculate_final_percentages(
        FRR=10,
        TFR=200,
        calcium_conc_mM=5
    )

    print("Final Formulation Composition and Lipid-to-Calcium Ratios")
    print("-" * 70)

    for component, value in results.items():
        if isinstance(value, str):
            print(f"{component:45}: {value}")
        elif "concentration" in component:
            print(f"{component:45}: {value:.3f} mM")
        elif "mol/mol" in component:
            print(f"{component:45}: {value:.2f} mol/mol")
        else:
            print(f"{component:45}: {value:.3f}%")
```
